# Efficacy of Drpitor1a, a Dynamin-Related Protein 1 inhibitor, in Pulmonary Arterial Hypertension

**DOI:** 10.1101/2023.12.21.572836

**Authors:** Danchen Wu, Ross D. Jansen-van Vuuren, Asish Dasgupta, Ruaa Al-Qazazi, Kuang-Hueih Chen, Ashley Martin, Jeffrey D. Mewburn, Elahe Alizadeh, Patricia D.A. Lima, Oliver Jones, Pierce Colpman, Nolan M. Breault, Isaac M. Emon, Luka Jedlovčnik, Yuan Yuan Zhao, Michael Wells, Gopinath Sutendra, Stephen L. Archer

**Affiliations:** Department of Medicine, Queen’s University, Kingston, ON K7L 3N6, Canada; Faculty of Chemistry and Chemical Technology, University of Ljubljana, Večna pot 113, 1000 Ljubljana, Slovenia; Queen’s Cardiopulmonary Unit (QCPU), Department of Medicine, Queen’s University, Kingston, ON K7L 3N6, Canada; Department of Medicine, University of Alberta, Edmonton, AB T6G 2R7, Canada; Partnerships and Innovation, Queen’s University, Kingston, ON K7K 3B5, Canada

**Keywords:** mitochondrial fission, super resolution STED microscopy, monocrotaline, pharmacokinetics, sexual dimorphism

## Abstract

**Rationale:** Dynamin-related protein 1 (Drp1), a large GTPase, mediates mitochondrial fission. Increased Drp1-mediated fission permits accelerated mitosis, contributing to hyperproliferation of pulmonary artery smooth muscle cells (PASMC), which characterizes pulmonary arterial hypertension (PAH). We developed a Drp1 inhibitor, Drpitor1a, and tested its ability to regress PAH.

**Objectives:** Assess Drpitor1a’s efficacy and toxicity in: a)normal and PAH human PASMC (hPASMC); b)normal rats versus rats with established monocrotaline (MCT)-induced PAH.

**Methods:** Drpitor1a’s effects on recombinant and endogenous Drp1-GTPase activity, mitochondrial fission, and cell proliferation were studied in hPASMCs (normal=3; PAH=5). Drpitor1a’s pharmacokinetics and tissue concentrations were measured (n=3 rats/sex). In a pilot study (n=3-4/sex/dose), Drpitor1a (1mg/kg/48-hours, intravenous) reduced adverse PA remodeling only in females. Consequently, we compared Drpitor1a to vehicle in normal (n=6 versus 8) and MCT-PAH (n=9 and 11) females, respectively. Drpitor1a treatment began 17-days post-MCT with echocardiography and cardiac catheterization performed 28-29 days post-MCT.

**Results:** Drpitor1a inhibited recombinant and endogenous Drp1 GTPase activity, which was increased in PAH hPASMC. Drpitor1a inhibited mitochondrial fission and proliferation and induced apoptosis, in PAH hPASMC but not normal hPASMC. Drpitor1a tissue levels were higher in female versus male RVs. In MCT-PAH females, Drpitor1a regressed PA obstruction, lowered pulmonary vascular resistance, and improved RV function, without hematologic, renal, or hepatic toxicity.

**Conclusions:** Drpitor1a inhibits Drp1 GTPase, reduces mitochondrial fission, and inhibits cell proliferation in PAH hPASMC. Drpitor1a caused no toxicity in MCT-PAH and had no significant effect on normal rats or hPASMCs. Drpitor1a is a potential PAH therapeutic which displays an interesting therapeutic sexual dimorphism.

## Introduction

Pulmonary arterial hypertension (PAH) is a cardiopulmonary syndrome, which has a ∼ 4/1 female predisposition(1). In PAH, obstruction, stiffening and vasospasm of small pulmonary arteries (PA) impairs functional capacity and leads to exertional dyspnea, syncope and ultimately death from right ventricular failure (RVF)(2). Obliteration of the small PAs is due partially to a cancer-like hyperproliferative, apoptosis-resistant, phenotype of pulmonary artery smooth muscle cells (PASMC), which increases pulmonary vascular resistance (PVR) and RV afterload(3). The RV in PAH is adversely affected by RV ischemia, adrenergic remodeling, fibrosis, mitochondrial metabolic changes, and inflammation(4, 5). Despite the many promising therapeutic targets identified in PAH(6), approved pharmacological treatments are primarily vasodilators. The prognosis for PAH remains poor, with >40% mortality at 5-years, suggesting new therapeutic paradigms are needed(7, 8).

Mitochondria not only synthesize ATP, but through their dynamic functions (fission, fusion, translocation and regulation of calcium homeostasis) regulate the cell cycle, and quality control processes such as mitophagy and apoptosis(9). Acquired disorders of mitochondrial dynamics in PAH constitute potential therapeutic targets(10). All cell types in the pulmonary vasculature in PAH, including PASMC(11), endothelial cells(12) and fibroblasts(13), have acquired abnormalities of mitochondrial metabolism (e.g. a shift toward uncoupled glycolysis) and/or dynamics (e.g. increased fission and depressed fusion), which in aggregate favour cell proliferation and inhibit apoptosis(9, 11, 14). There are no approved mitochondrial-targeted therapies for PAH. One promising therapeutic target is dynamin-related protein 1 (Drp1), a large GTPase that mediates mitochondrial fission(15). When activated, Drp1 translocates to the outer mitochondrial membrane (OMM) where it forms a multimeric protein complex with one or more of its four binding partners(16). The resulting ring-like structure uses Drp1’s GTPase function to constrict and divide the mitochondrion. During mitosis, there is an obligatory coordination between mitochondrial mitotic fission and nuclear division, which ensures equal distribution of mitochondria and mitochondrial DNA to the daughter cells(9).

Since Drp1’s GTPase activity is essential for fission(15, 17), this domain is a logical pharmacologic target. Expression of Drp1 and its adapter proteins, mitochondrial dynamics protein of 49 and 51 kDa (MiD49 and MiD51), are increased in human and experimental PAH(18, 19). Inhibiting Drp1, using the Drp1 GTPase inhibitor, mdivi-1, or small inhibitory RNA (siDrp1), causes G2/M phase cell cycle arrest and apoptosis in PAH hPASMC(19, 20). The importance of fission is further highlighted by the observation that downregulation of MiD49 and MiD51 prevents fission, and also causes cell cycle arrest, and regresses PAH in PAH hPASMC and rodent PAH models(18).

Because of concerns regarding the specificity of mdivi-1, we developed a more potent and specific small molecule inhibitor, Drpitor1a, as a potential therapeutic agent. Drpitor1a is an ellipticine- like compound identified in a chemical library based on its predicted ability to bind Drp1’s GTPase domain. Drpitor1a specifically inhibits Drp1 GTPase in cancer cells and is 50-fold more potent than mdivi-1 in inhibiting fission(21). Drpitor1a regresses lung cancer in a xenotransplantation murine model and protects the RV from ischemia-reperfusion injury(22).

Here, we show that Drpitor1a inhibits Drp1’s GTPase which inhibits fission without preventing Drp1’s translocation to mitochondria. We established the pharmacokinetics of Drpitor1a and showed greater drug accumulation in the RV of females. *Ex vivo* Drpitor1a slows PAH hPASMC proliferation, increases apoptosis and *in vivo* regresses MCT-PAH. Reassuringly, Drpitor1a has minimal effects on fission in normal cells and elicits no toxicity in MCT-PAH rats or normal rats. We observed a sexual dimorphism, such that Drpitor1a is more effective in female than male MCT- PAH rats. Drp1 is a promising therapeutic target in PAH and Drpitor1a a potential therapeutic molecule.

## Methods

For detailed materials and methods used in this study, please see Online Data Supplement.

### 1. *In vitro* studies Cell Culture

#### 1.1. Compound synthesis

A modified reaction scheme of Drpitor1a synthesis is shown in **Figure E1**.

#### 1.2. Cell Culture

hPASMC from normal donors (n=3; 67% female, mean age: 50.7 years) and patients with PAH (n=5; 40% female, mean age: 41.2 years) were maintained as reported(23).

#### 1.3. GTPase assay

GTPase activity of both recombinant Drp1 and immunoprecipitated Drp1 from PAH hPASMC, was quantified using a colorimetric assay that detects phosphate (ab270553, Abcam, Toronto, ON, Canada).

#### 1.4. Mitochondrial morphology analysis and measurement of endogenous Drp1

Normal or PAH hPASMC were treated with Drpitor1a (0.1 μM) or DMSO for 4-5 hours. Mitochondria were stained in live cells using the potentiometric dye, tetramethylrhodamine methyl ester (TMRM, 20nM), and imaged as reported(23). Mitochondrial fragmentation was quantified using two published metrics: mitochondrial fragmentation count (MFC) and a machine-learning based algorithm(22).

#### 1.5. Immunofluorescence imaging of endogenous Drp1

Normal or PAH hPASMC were treated with Drpitor1a (0.1 μM) or DMSO for 5 hours and stained with MitoTracker Deep Red (MTDR). Cells were then fixed and total Drp1 on mitochondria was imaged using a confocal microscope.

#### 1.6. Immunoblots

Methodology and reagents are reported in the Online Data Supplement.

#### 1.7. Cell proliferation

hPASMC proliferation was measured 48 hours post-Drpitor1a treatment (1 μM).

#### 1.8. Cell apoptosis

hPASMC apoptosis was determined 48 hours post-Drpitor1a treatment (5 μM) by the Annexin V+ propidium iodide (PI) method.

## 2. *In vivo* studies

The animal experiment protocol was approved by the University Animal Care Committee of Queen’s University (protocol 2021-2112).

### 2.1. Pharmacokinetics analysis

A single dose of Drpitor1a (1mg/kg or 5mg/kg) was administered IV or PO (n=3/sex). Pharmacokinetic analysis on blood samples was performed at the Platform of Biopharmacy, University of Montreal using an LC/MS/MS AB/SCIEX 4000 QTRAP.

### 2.2. Tissue Drpitor1a concentration analysis

RV and lung samples were snap frozen and stored at -80 °C. Tissue concentrations of Drpitor1a was measured using HPLC coupled with mass spectrometry.

### 2.3. Efficacy of Drpitor1a on MCT-PAH

#### 2.3.1. Sex and sample size

##### Pilot study

The pilot study was designed to assess feasibility and tolerability of Drpitor1a and observe potential efficacy signals (defined as a trend to reduce adverse remodeling of small PAs or regress RV hypertrophy in MCT-PAH). Three rats/sex were used in the MCT+NS groups and 4 rats/sex were used in the two MCT+Drpitor1a groups (0.1 mg/kg and 1.0 mg/kg IV every 48 hours) (**Figure 4A**).

##### Confirmation study

Because Drpitor1a lacked a benefit trend in male rats in the pilot study, only female rats were used in the confirmation study. We compared Drpitor1a to vehicle in normal females (n=6 vs 8) and MCT-PAH females (n=9 and 11), respectively. Drpitor1a treatment (1.0 mg/kg every 48 hours) began 17-days post-MCT with echocardiography and cardiac catheterization performed 28-29 days post-MCT.

#### 2.3.2. **MCT-PAH**

See Online Data Supplement.

#### 2.3.3. Left jugular vein indwelling catheter implantation

See Online Data Supplement.

#### 2.3.4. Drpitor1a treatment of female rats with established MCT-PAH

Drpitor1a (1 mg/kg) or vehicle (5% DMSO) was administered IV every 48 hours from day 17 to day 27.

#### 2.3.5. Echocardiography

Doppler, 2-dimensional (2-D), and M-mode ultrasound was performed on days 14 and 28 post-MCT/PBS injection, as described (24). See Online Data Supplement for details and measured parameters.

#### 2.3.6 **Hemodynamics-left and right heart catheterization (RHC)**

Pulmonary hemodynamics were measured on day 29 by RHC delivered via a closed-chest approach, as described (18). RV systolic pressure (RVSP) and cardiac output (CO) were measured.

#### 2.3.7. **Blood collection**

Whole blood was collected via cardiac puncture on day 29 after RHC.

### Statistical analyses

Data are presented as mean ± standard error of the mean (SEM). Normality of the data sets was determined using the Shapiro-Wilk test. A Student t-test or Mann-Whitney U test was used to compare the mean or median value between two groups, as appropriate. One-way or two-way ANOVA with post hoc analyses were used, as appropriate. A P-value of < 0.05 was considered statistically significant. Analyses were performed using GraphPad-Prism 9 software (San Diego, CA, USA).

## Results

### Drpitor1a is a Drp1 GTPase inhibitor that prevents fission which inhibits accelerated proliferation and promotes apoptosis in PAH hPASMC

The GTPase activity of recombinant human Drp1 is inhibited by Drpitor1a (P=0.021, **Figure 1A**). The GTPase activity of endogenous Drp1 is significantly increased in PAH hPASMC versus control hPASMC (P=0.029, **Figure 1B**, **Figure E2**) and is also inhibited by Drpitor1a (P <0.001, **Figure 1C**). Drpitor1a inhibited mitochondrial fission in PAH hPASMC, decreasing the MFC (P<0.001, **Figure 1D**) and increasing the percentage area of filamentous mitochondria (P<0.001, **Figure 1E**). In normal hPASMC, Drpitor1a did not alter mitochondrial morphology (**Figure 1F**). Despite inhibiting fission, Drpitor1a increased the accumulation of endogenous Drp1 on mitochondria in PAH hPASMC, as measured by confocal microscopy (**Figure 1G**). Drpitor1a did not alter total Drp1 expression in PAH hPASMC (Figure E3), as measured by immunoblotting. Drpitor1a reduced cell proliferation rates in PAH hPASMC (P=0.05, **Figure 1H**) without significantly reducing the much lower proliferation rate of control hPASMC. Drpitor1a increased unstimulated apoptosis in two out of three PAH hPASMC cell lines, with an accompanying significant increase in the expression of the apoptosis mediator Bak and suppression of the anti-apoptosis mediator, Bcl-2 (**Figure 1I-J**, Figure E4).

**Figure 1.**
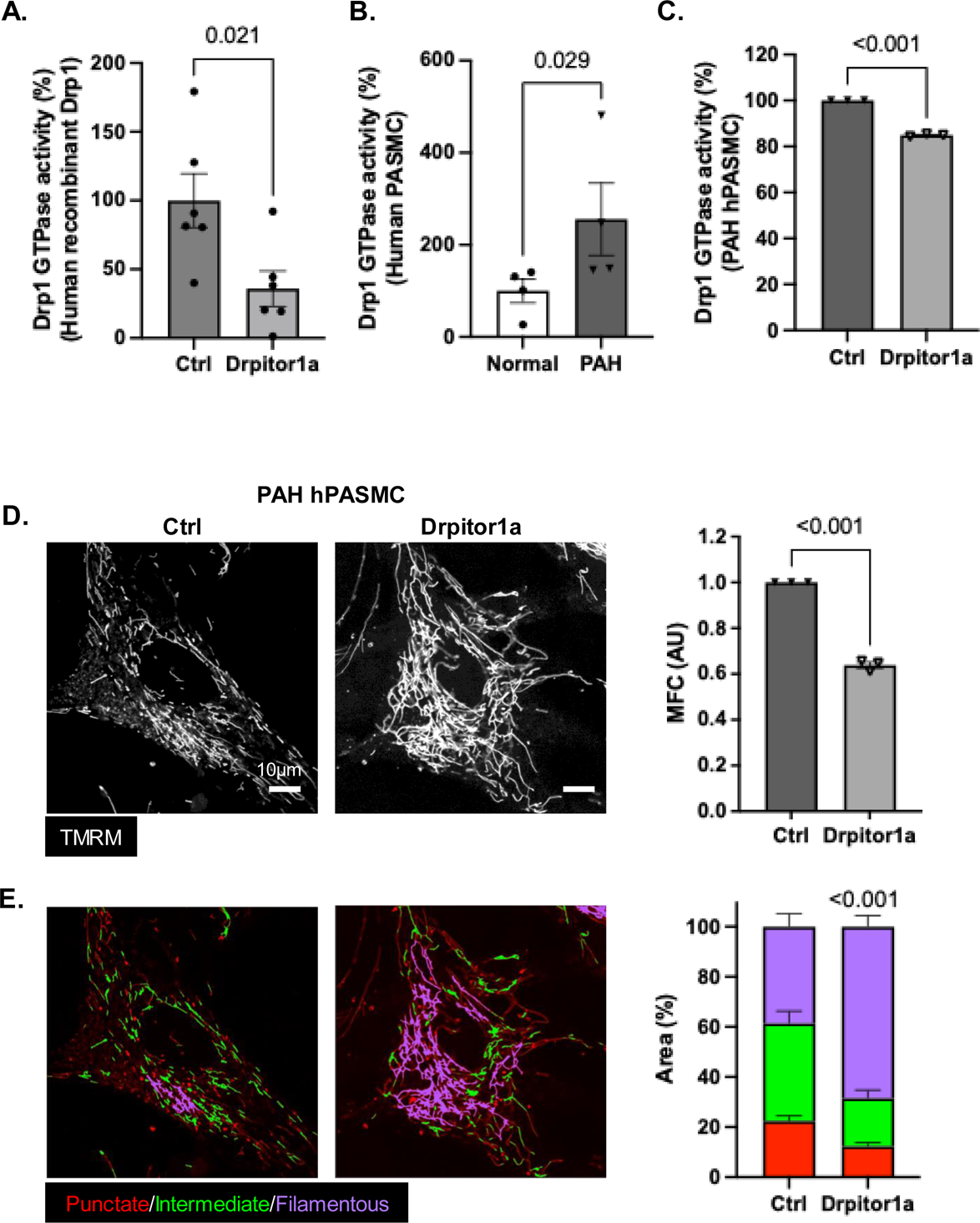
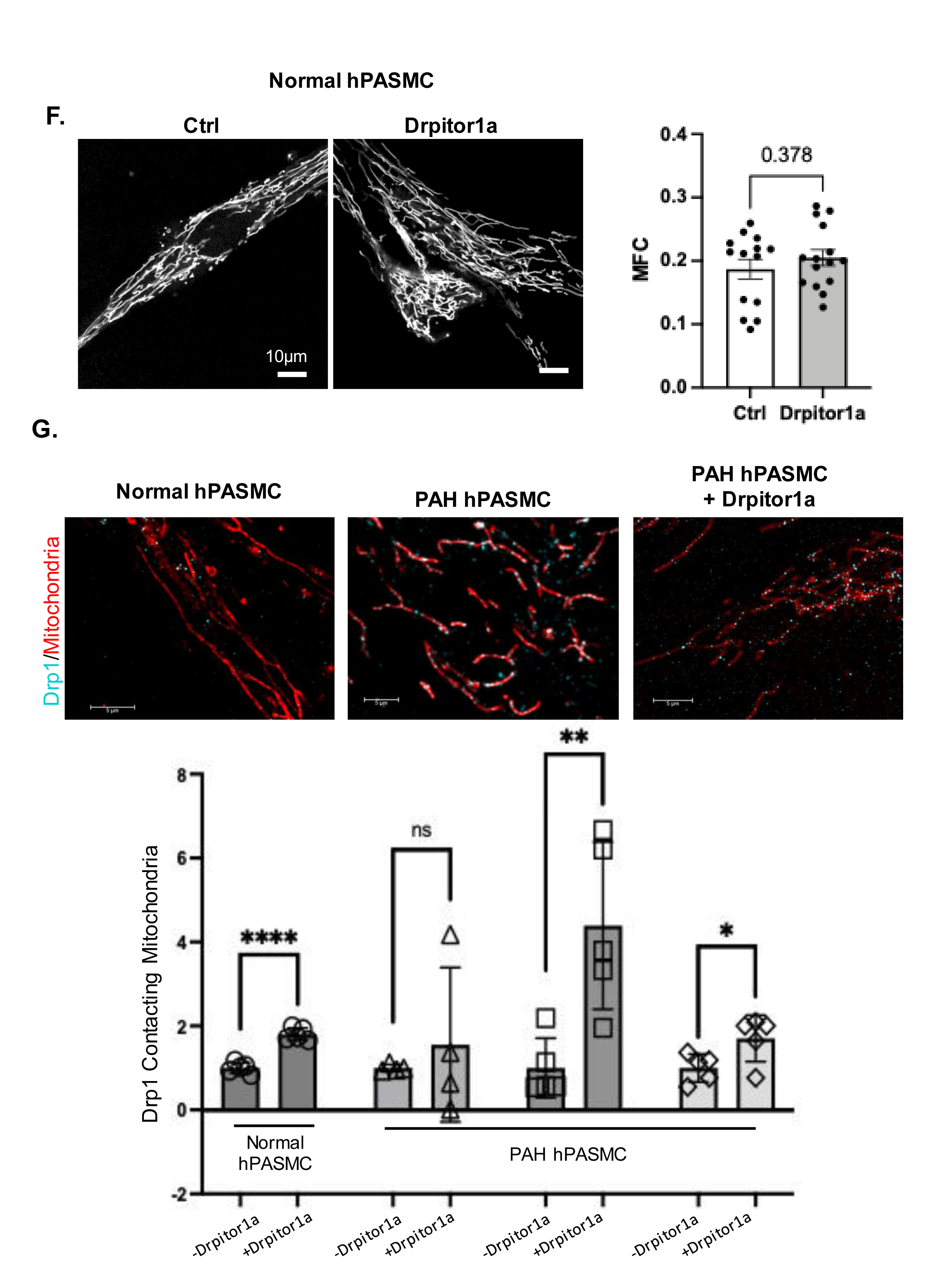
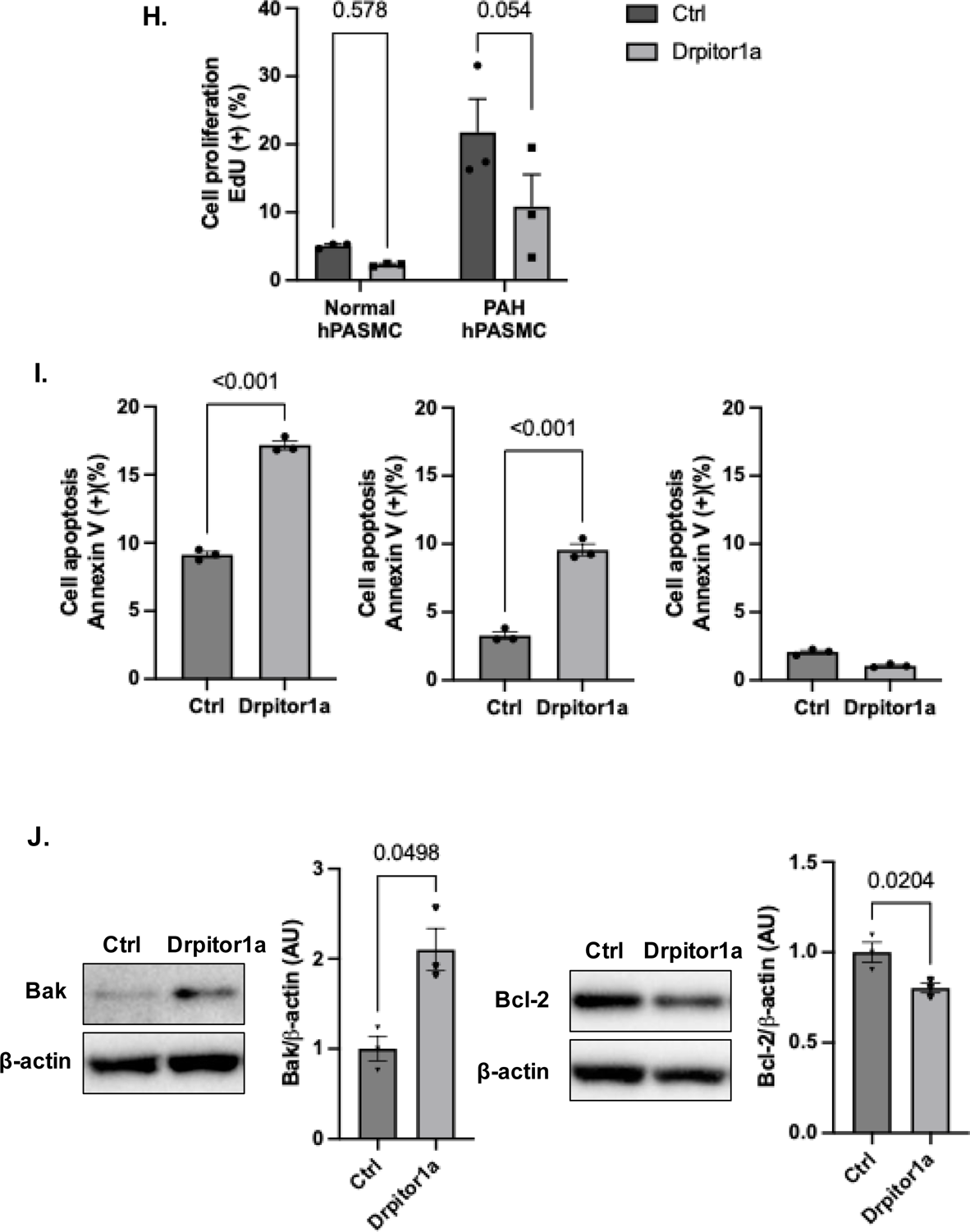
Drp1 GTPase activity is increased in human pulmonary artery smooth muscle cells (hPASMC) from patients with PAH, and Drpitor1a reverses the PAH phenotype. **A. Drpitor1a inhibits the GTPase activity of human recombinant Drp1** (n=6). **B. Drp1 GTPase activity is significantly increased in PAH hPASMC compared to normal hPASMC** (n=4 for normal and n=4 for PAH). **C. Drp1 GTPase activity is significantly inhibited by Drpitor1a in PAH hPASMC** (n=3). **D. Representative confocal microscopy images of PAH hPASMC showing mitochondrial fission is significantly inhibited by Drpitor1a. Mitochondria were stained with 20nM tetramethylrhodamine, methyl ester (TMRM). Mitochondrial fission was quantified by mitochondrial fragmentation count (MFC)** (n=3). **E.** Color-coded confocal microscopy images showing the percentage area of filamentous mitochondria is significantly increased by Drpitor1a. Red=punctate mitochondria; green= intermediate mitochondria; purple=filamentous mitochondria. F. Drpitor1a does not change mitochondrial morphology in normal hPASMC. G. Drpitor1a does not inhibit the translocation of exogenous Drp1 to mitochondria. Endogenous Drp1 (cyan) accumulates on TMRM loaded mitochondria (red) more in PAH PASMC following treatment with Drpitor1. **, **** p<0.01 and 0.0001, respectively. **H.** Cell proliferation of PAH hPASMC is significantly inhibited by Drpitor1a, while Drpitor1a does not significantly inhibit cell proliferation in normal hPASMC. I. Apoptosis of PAH hPASMC is significantly induced by Drpitor1a in 2 out of 3 cell lines. J. Apoptosis of PAH hPASMC is significantly induced by Drpitor1a. There is increased expression of the pro-apoptotic protein, Bak, and decreased expression of the anti- apoptotic protein, Bcl-2.

### Pharmacokinetics, bioavailability, and tissue concentration of Drpitor1a in normal rats

Drpitor1a was administered at 1 mg/kg or 5 mg/kg in all *in vivo* studies (protocol in **Figure 2A**, **Table 1, Table E2**). The half-life of Drpitor1a (1 mg/kg, IV) is shorter in females (3.4±0.7 hours) than males (6.5±0.8 hours) and increases with high dose (5 mg/kg) in both sexes (**Table 1**, **Figure 2B-C**). The oral bioavailability of Drpitor1a at the 1mg/kg dose is lower in females than males (female 12.6% versus male 26.0%); but this sex difference disappears at the 5mg/kg dose (**Table 1**).

**Figure 2.**
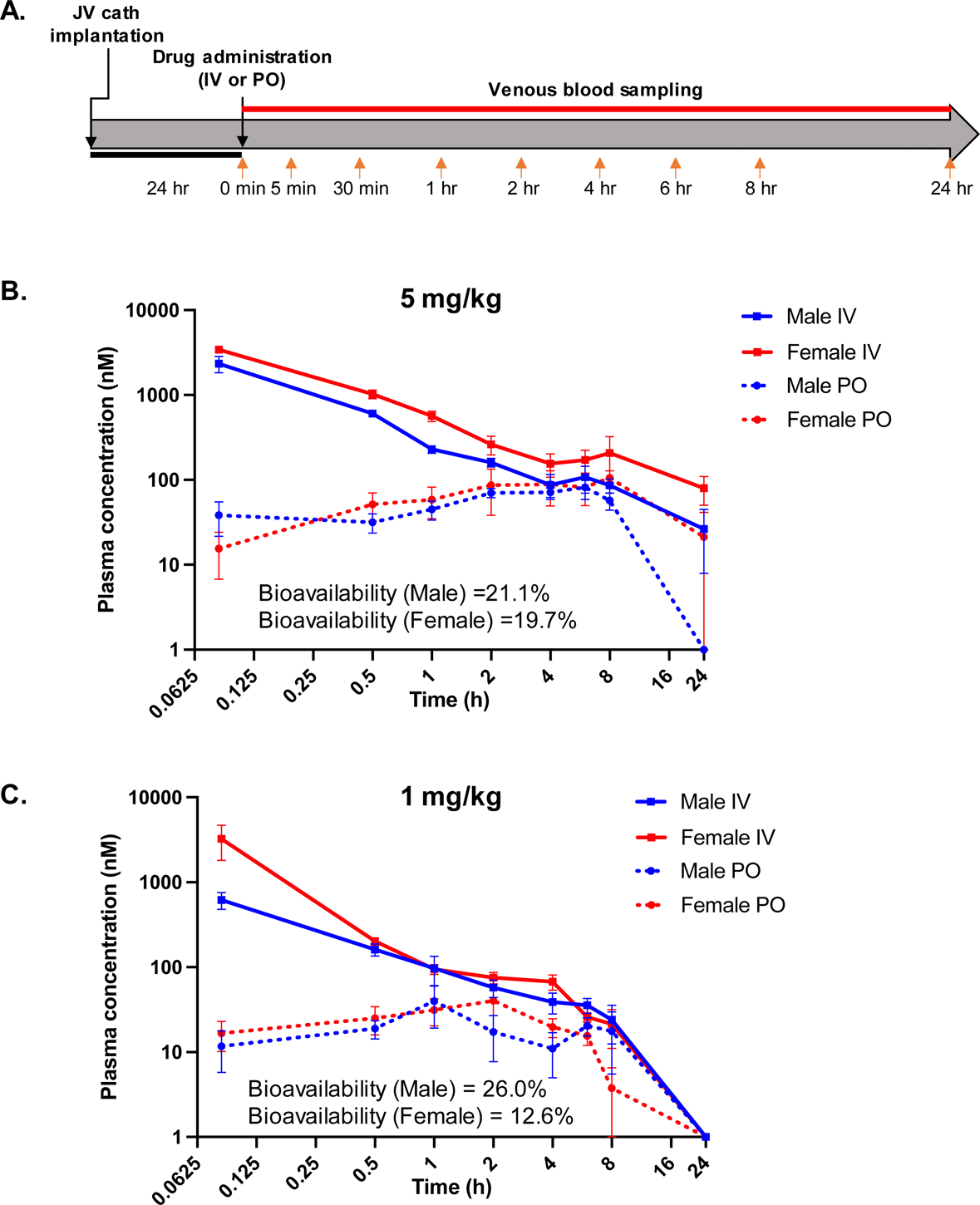
**Plasma concentration of Drpitor1a in the pharmacokinetics (PK) study.** A. Timeline of the PK study in rats. B. Plasma concentration of Drpitor1a in rats after a single dose administration (5mg/kg) (n=3/sex/route). C. Plasma concentration of Drpitor1a in rats after a single dose administration (1mg/kg) (n=3/sex/route).

**Table 1.** Average ± Standard deviation of the pharmacokinetic parameters in the rats receiving intravenous administration of Drpitor1a.

| **Intravenous administration** |  |  |  |  |
| --- | --- | --- | --- | --- |
| **Parameters** | **Male - 1mg/kg** | **Female - 1mg/kg** | **Male - 5mg/kg** | **Female - 5mg/kg** |
| AUC<sub>0-t</sub> (nM*h) | 552.5 ± 223.3 | 1288.2 ± 702.1 | 2496.9 ± 1288.2 | 5306.6 ± 2790.9 |
| AUC<sub>0-∞</sub> (nM*h) | 844.3 ± 310.9 | 1433.8 ± 754.6 | 3211.9 ± 1769.4 | 8577.4 ± 4719.0 |
| MRT (h) <sup>(a)</sup> | 7.2 ± 0.8 | 2.4 ± 0.8 | 10.4 ± 6.5 | 27.5 ± 27.8 |
| kel (h) | 0.1 ± 0.01 | 0.2 ± 0.04 | 0.08 ± 0.03 | 0.05 ± 0.03 |
| t<sub>1/2</sub> terminal (h) | 6.5 ± 0.8 | 3.4 ± 0.7 | 9.9 ± 5.3 | 24.3 ± 21.4 |
| T<sub>max</sub> (h) | 0.08 ± 0 | 0.08 ± 0 | 0.08 ± 0 | 0.08 ± 0 |
| C<sub>max</sub> (nM) | 617.7 ± 236.5 | 3250.4 ± 2499.6 | 2332.3 ± 884.2 | 3425.0 ± 535.1 |
| CL<sub>p</sub> (mL/min) | 23.9 ± 11.2 | 13.1 ± 5.3 | 39.8 ± 22.2 | 13.7 ± 10.8 |
| V<sub>d</sub> (L) | 10.2 ± 4.3 | 1.9 ± 1.2 | 20.3 ± 6.0 | 14.3 ± 8.4 |
| % Bioavailability<br>(calculated with<br>corresponding IV<br>AUC 0-t) | 26.0 | 12.6 | 21.1 | 19.7 |

(a) MRT = time for elimination of 63.2% of the dose. AUC: Area Under the Concentration-Time Curve MRT: Mean Residence Time kel: elimination rate constant t<sub>1/2</sub>: terminal half-life T<sub>max</sub>: the time it takes for Drpitor1a to reach its maximum concentration in the bloodstream after administration. C<sub>max</sub>: the maximum concentration of Drpitor1a in the bloodstream after it has been administered. CL<sub>p</sub>: the clearance of Drpitor1a from the plasma V<sub>d</sub>: the volume of distribution

### Lung concentrations of Drpitor1a

The 1 mg/kg dose used in our studies for therapy yielded higher lung Drpitor1a levels both in IV (0.910±1.473 nmole/mg) and PO (0.013 ± 0.004 nmole/mg) in females compared to males, although the difference between sexes was not statistically significant (P > 0.999 and P=0.2444 for PO and IV respectively). Oral Drpitor1a (5 mg/kg) yields a higher lung concentration in females (0.021 ± 0.030 nmole/mg) vs males (0.002±0.001 nmole/mg) but this is not statistically significant (P > 0.999). When given IV, the 5 mg/kg dose yields much higher lung concentrations, but levels are similar in females (0.444±0.581 nmole/mg) and males (0.441 ± 0.015nmole/mg) (P > 0.999) (**Figure 3A**).

**Figure 3.**
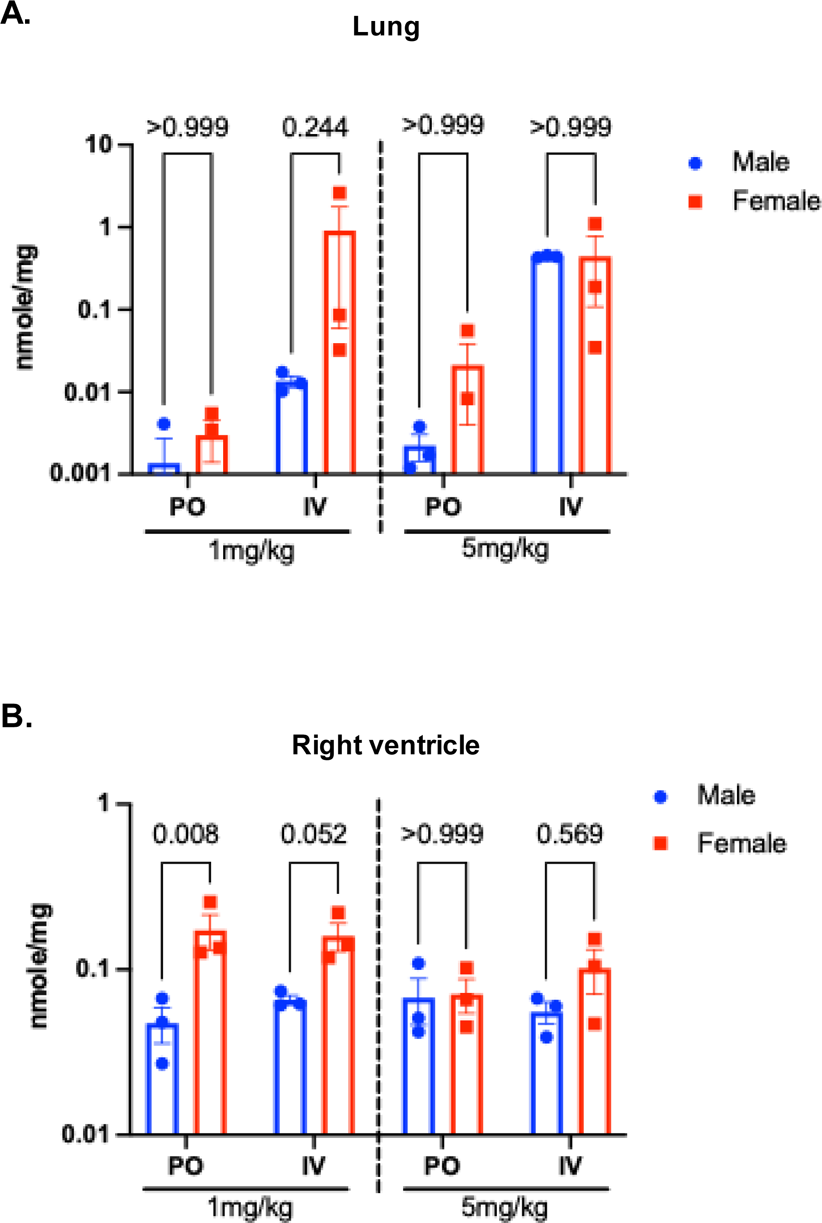
**Tissue concentration of Drpitor1a in rats.** A. The concentration of Drpitor1a in lung tissue 24 hours after a single dose of Drpitor1a (n=3/sex/dose/route). B. The concentration of Drpitor1a in right ventricle tissue 24 hours after a single dose of Drpitor1a (n=3/sex/dose/route).

### RV concentrations of Drpitor1a

Intravenously, 1 mg/kg Drpitor1a also yields higher RV drug concentrations in females versus males (0.160±0.054 versus 0.066±0.007 nmole/mg; P=0.052, **Figure 3B**). Twenty-four hours after a single, oral, 1 mg/kg dose RV Drpitor1a levels are lower than IV but still are significantly higher in females (0.173±0.072nmole/mg) than males (0.047±0.020 nmole/mg) (P=0.008). The 5 mg/kg dose IV yields slightly higher concentrations in females (0.102±0.053 nmole/mg) versus males (0.055±0.015 nmole/mg) (P=0.5686) whilst the 5 mg/kg oral dose yields similar RV concentrations in females (0.071±0.029 nmole/mg) and males (0.067±0.036 nmole/mg) (P>0.999) (**Figure 3B**, Figure E5).

### Therapeutic efficacy of Drpitor1a in the rat MCT-PAH model

#### Pilot study

Drpitor1a (1 mg/kg IV every 48 hours) was well tolerated (no deaths or weight loss) and showed a statistically insignificant trend to decrease medial thickness of the small PAs in female (but no trend in male) MCT rats (P=0.141, **Figure 4B**). Drpitor1a also showed a statistically insignificant trend to reduce RV hypertrophy only in female MCT-PAH rats (P=0.449, **Figure 4C**). Consequently, only female rats were used in the confirmation study, which was adequately powered to detect changes in histology and hemodynamics.

#### Confirmation study

The design of the confirmation study is shown in **Figure 5A**. Ultrasound on day 14 post-MCT quantified pulmonary hypertension (PH), evident as decreased PAAT and adverse changes to RV structure and function (evident as increases in RV free wall (RVFW) thickness and decreases in RVFW thickening, TAPSE and cardiac index (CI)) (P<0.001 for all parameters, **Figure 5B-F**). At randomization, there was no significant difference in these ultrasound parameters between MCT rats, which were randomly assigned to receive normal saline (NS) or Drpitor1a (**Figure E6**).

**Figure 4.**
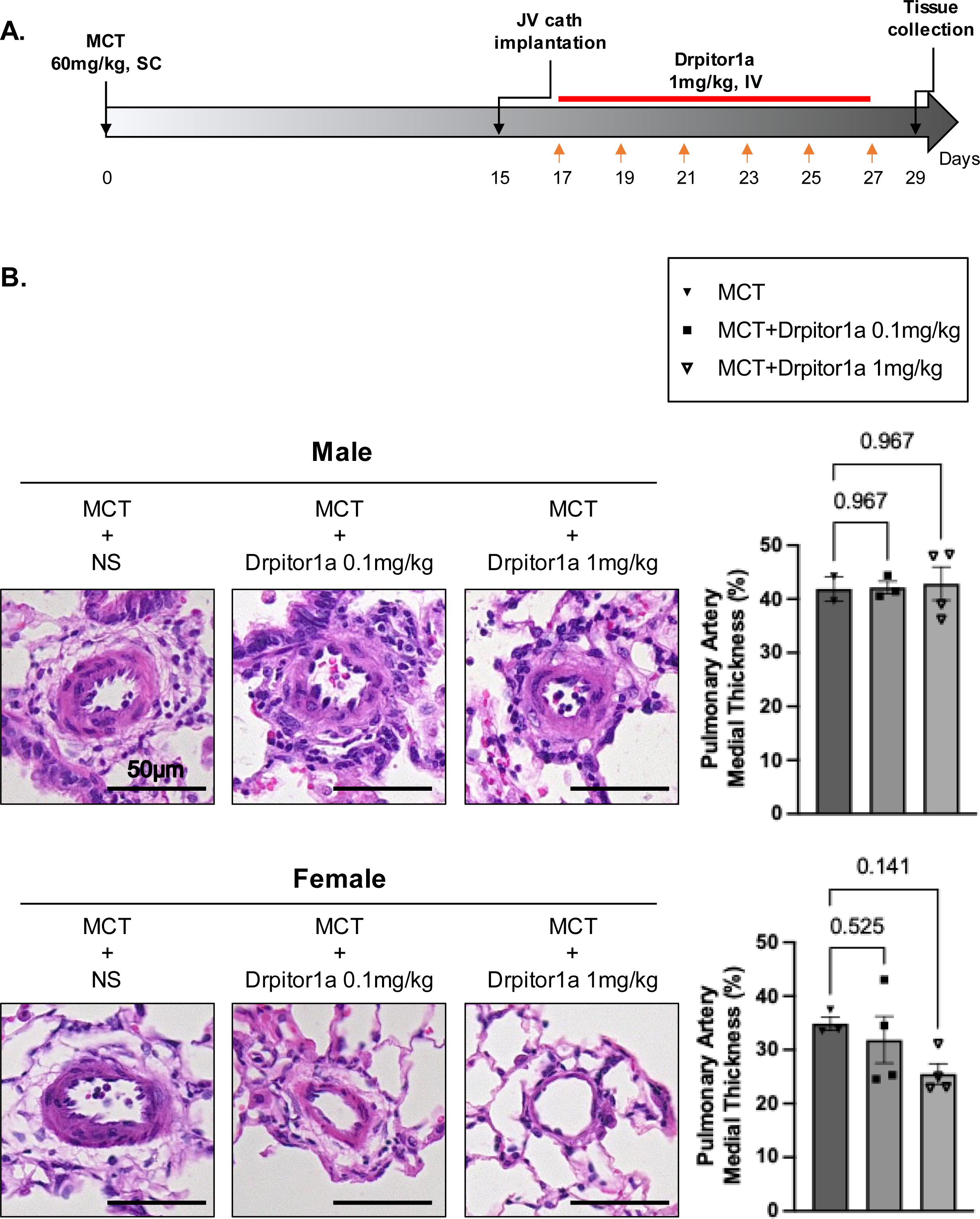
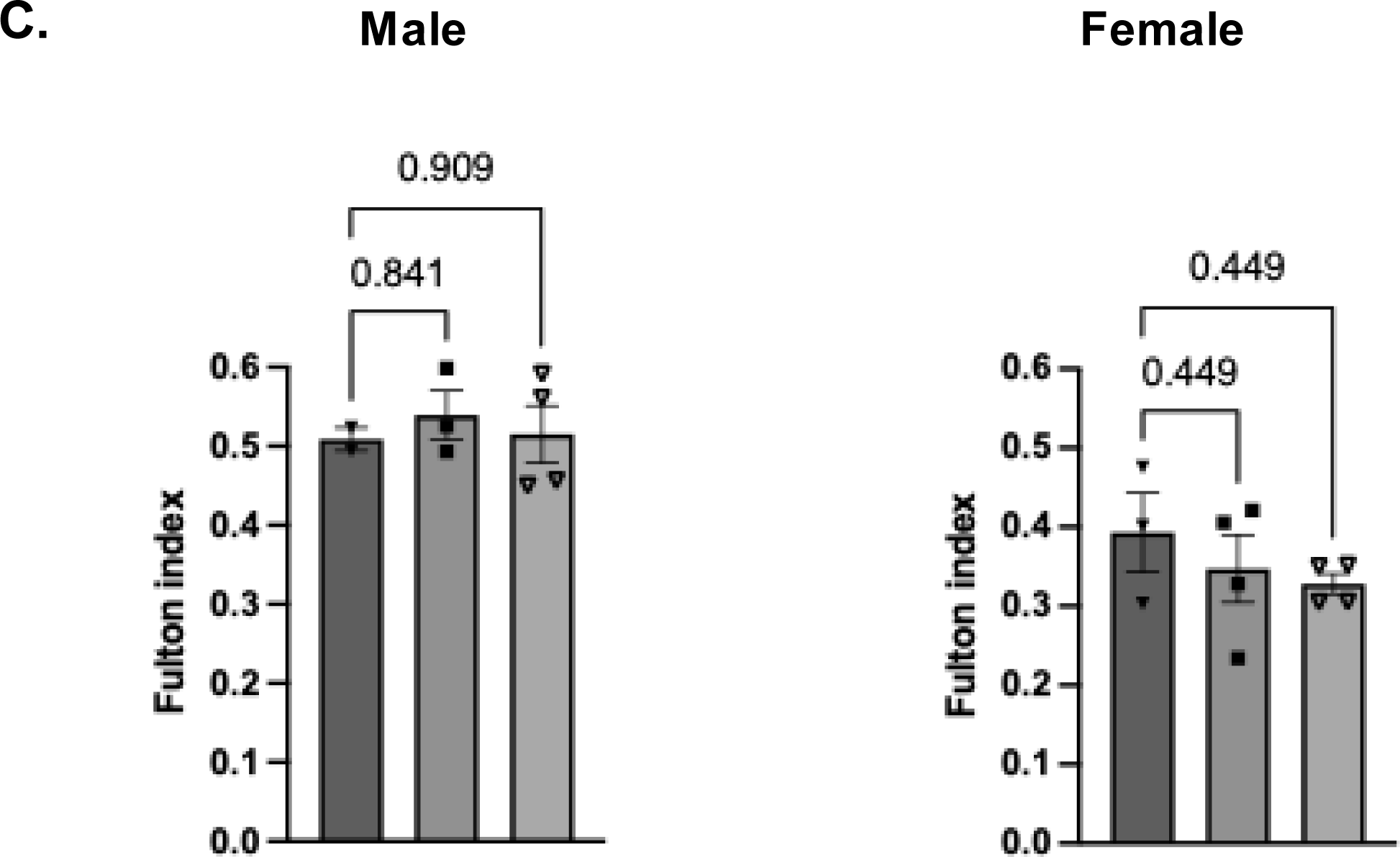
**Drpitor1a regressed PAH in female MCT-PAH rats only.** A. Study design for the pilot study. B. Pulmonary vascular histology showing Drpitor1a reduced %medial thickness of small PA in female MCT-PAH rats. H&E staining. C. Fulton index showing Drpitor1a tended to regress RV hypertrophy in female but not male MCT- PAH rats. Male: n=2 for MCT+NS, n=3 for MCT+Drpitor1a 0.1mg/kg, n=4 for MCT+Drpitor1a 1mg/kg. Female: n=3 for MCT+NS, n=4 for MCT+Drpitor1a 0.1mg/kg, n=4 for MCT+Drpitor1a 1mg/kg.

**Figure 5.**
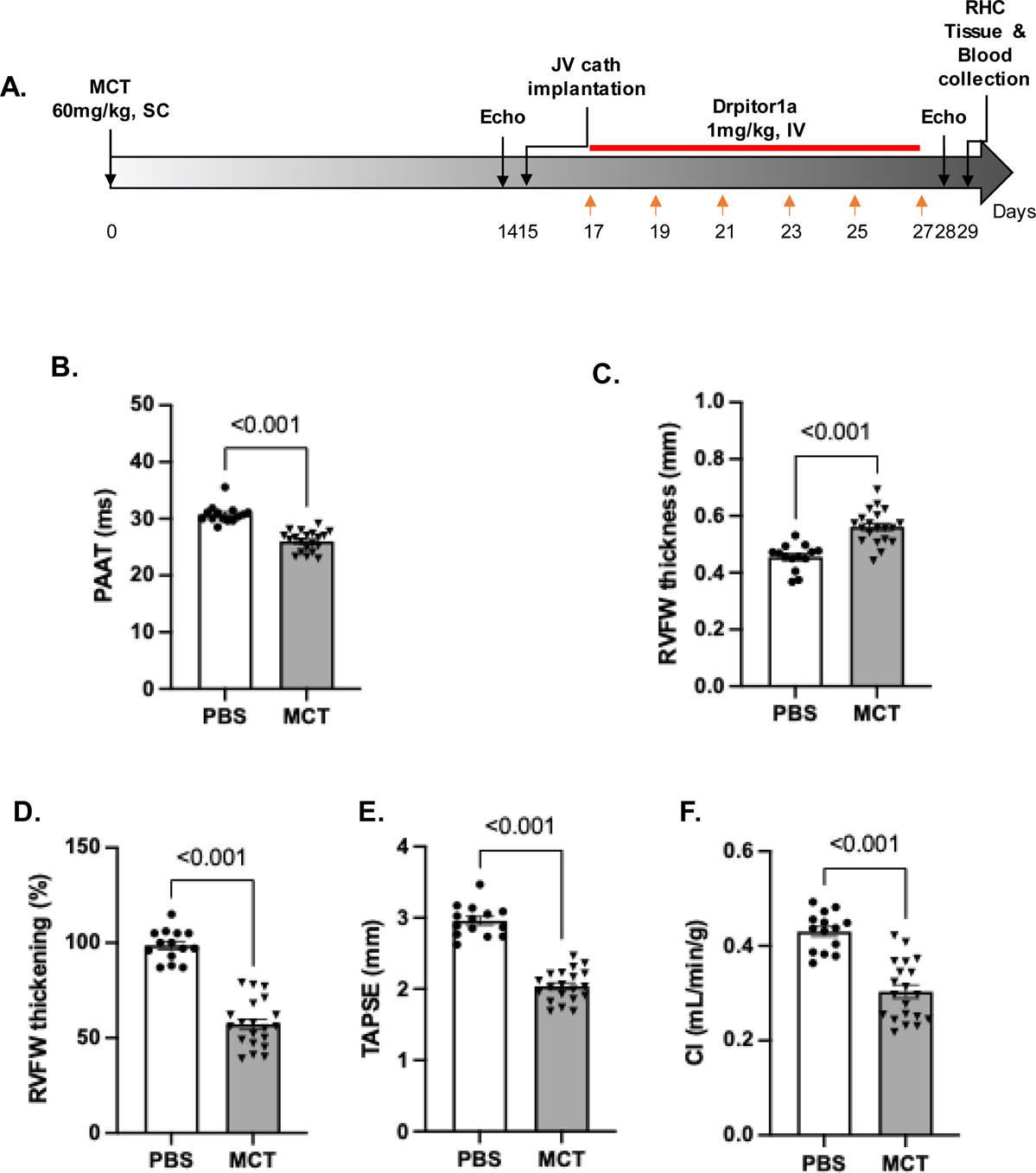
Development of pulmonary hypertension and RV dysfunction 14 days post-MCT (n=14 for PBS and n=20 for MCT). A. Timeline for the confirmation study. B. MCT rats have shortened pulmonary artery acceleration time (PAAT) compared to the PBS rats. C. MCT rats have increased right ventricular free wall thickness (RVFWT) compared to the PBS rats. D. MCT rats have decreased right ventricular free wall thickening (RVFWT%) compared to the PBS rats. E. MCT rats have decreased tricuspid annular plane systolic excursion (TAPSE) compared to the PBS rats. F. MCT rats have lower cardiac index (CI) compared to the PBS rats.

### Drpitor1a reversed adverse pulmonary vascular remodeling and RV hypertrophy and decreased PH severity in MCT-PAH

The percent medial thickness of the small PAs was higher in MCT+NS rats compared to the PBS+NS rats (P<0.001), and this adverse vascular remodeling was decreased by Drpitor1a treatment (P<0.001, **Figure 6A**). Furthermore, the cross-sectional RV cardiomyocyte area, a measure of RV hypertrophy, which was significantly increased in the MCT+NS rats (P<0.001), was decreased by Drpitor1a (P<0.001, **Figure 6B**). There was no change in PA medial thickness and RV cardiomyocyte area between the PBS+NS and PBS+Drpitor1a groups (P=0.996 and 0.653, respectively, **Figure 6A-B**).

**Figure 6.**
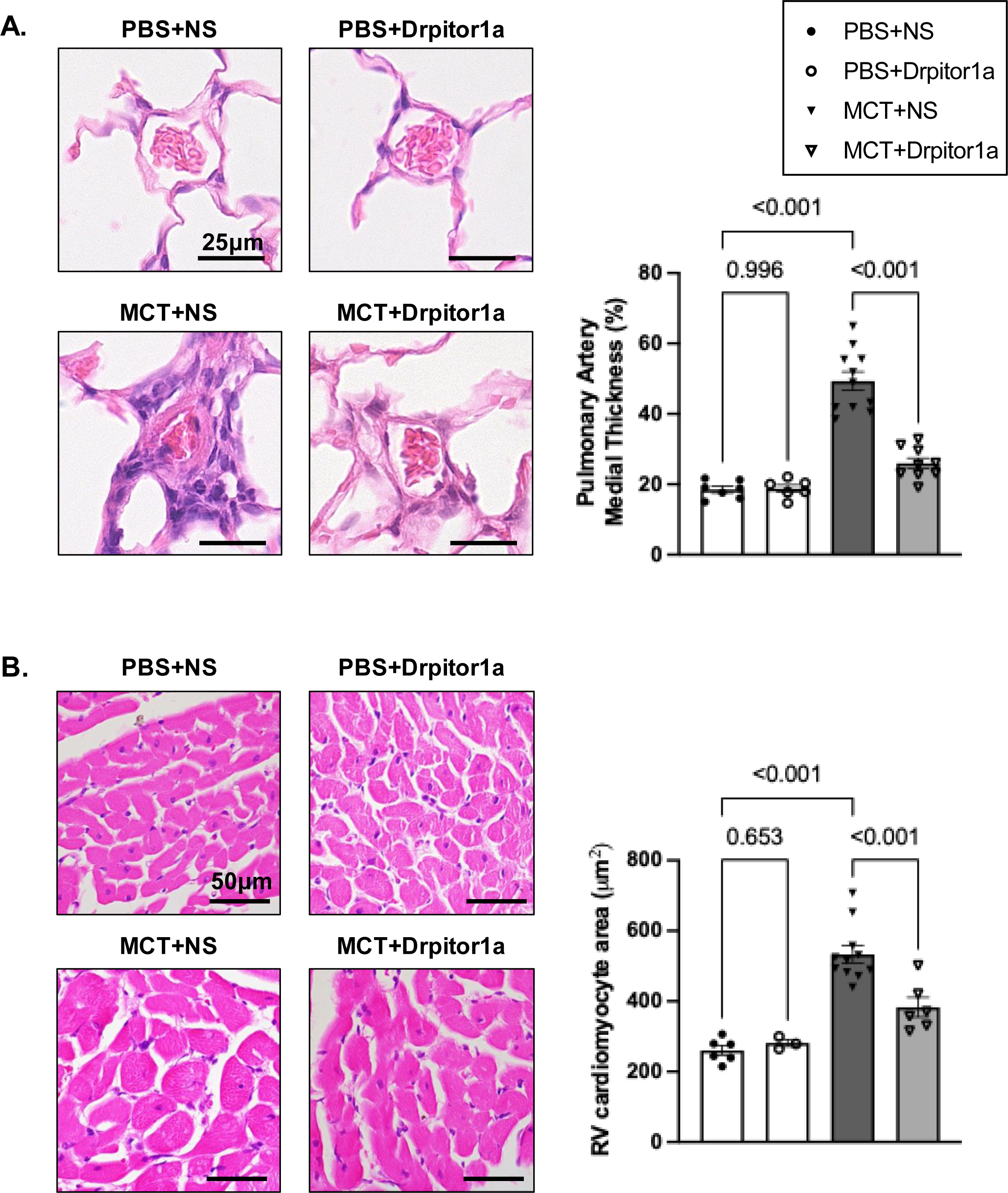
Drpitor1a inhibited adverse pulmonary vascular remodeling and RV hypertrophy in MCT-PAH rats. (n=14 for PBS, n=20 for MCT). A. Drpitor1a reversed pulmonary vascular remodeling as measured by reduced % medial thickness of small pulmonary artery in MCT-PAH rats. B. Drpitor1a reversed RV hypertrophy as measured by RV cardiomyocyte cross-sectional area in MCT-PAH rats.

At the end of the study (29 days), MCT+Drpitor1a rats had less severe pulmonary hypertension than the MCT+NS rats, as evidenced by significantly longer PAAT (P=0.020) and lower PVR (P=0.044, **Figure 7A-B**). Drpitor1a also improved RV function and reduced RVH, as shown by increased RVFW thickening (P=0.042), TAPSE (P=0.010), and CI (P<0.001); and reduced RVFW thickness (P=0.048) and Fulton index (P=0.015), respectively (**Figure 7A-C**, Figure E7). Measures of RV coupling, TAPSE/mean pulmonary artery pressure (mPAP) (P=0.618) and RVFW thickening/mPAP (P>0.999), were not significantly improved in the MCT+Drpitor1a group. Thus Drpitor1a improved RV function without enhancing RV-PA coupling (**Figure 7D**).

**Figure 7.**
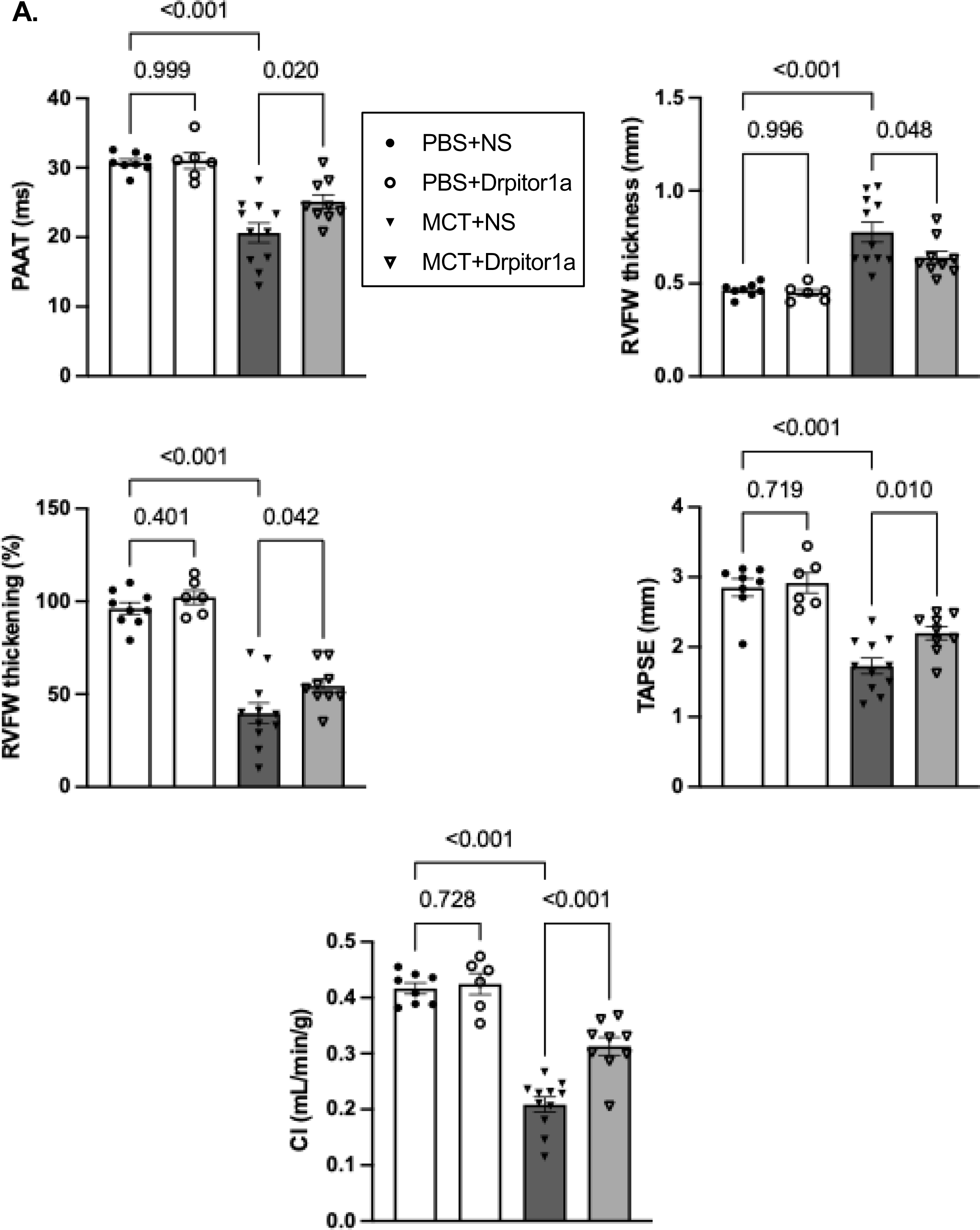
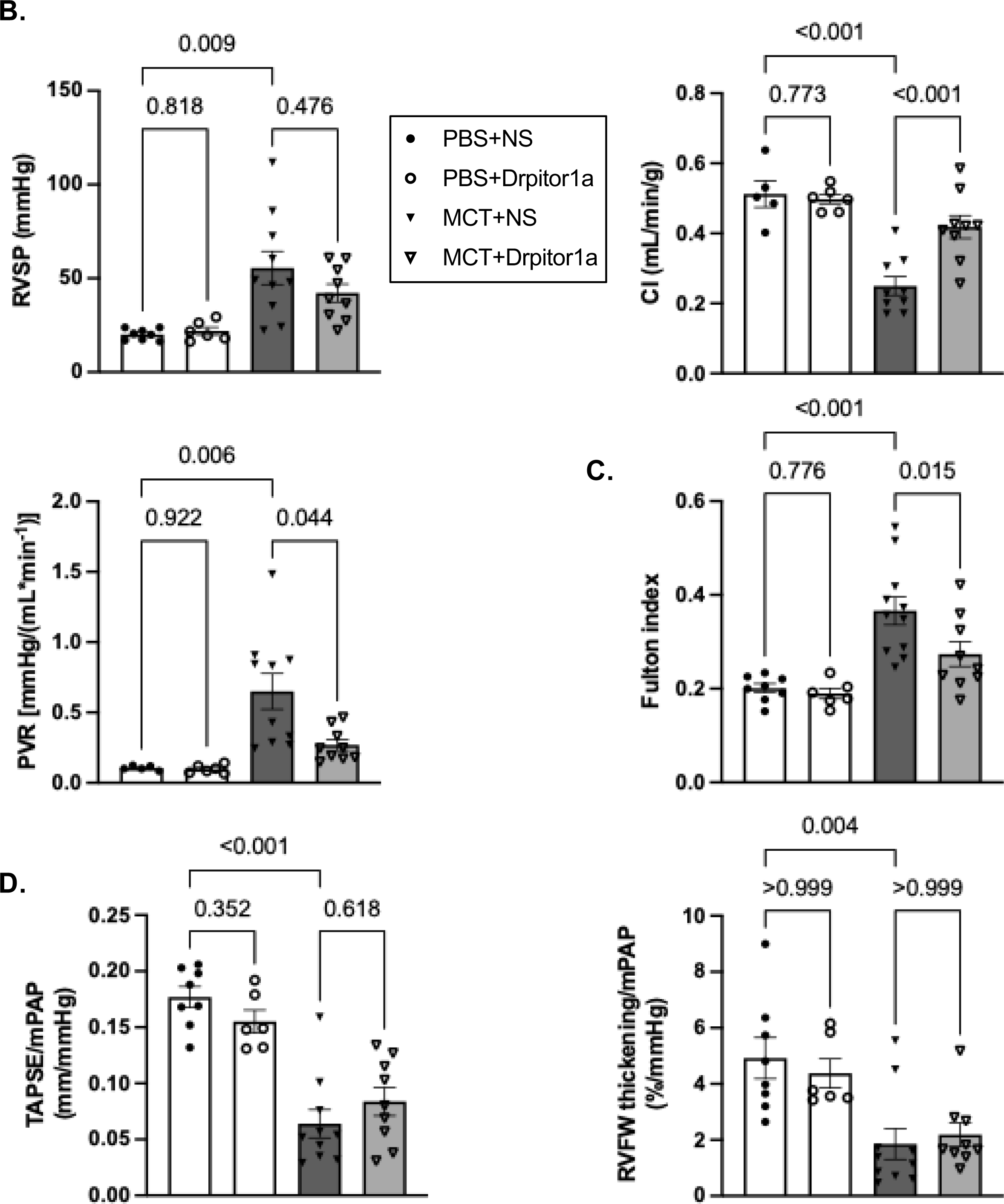
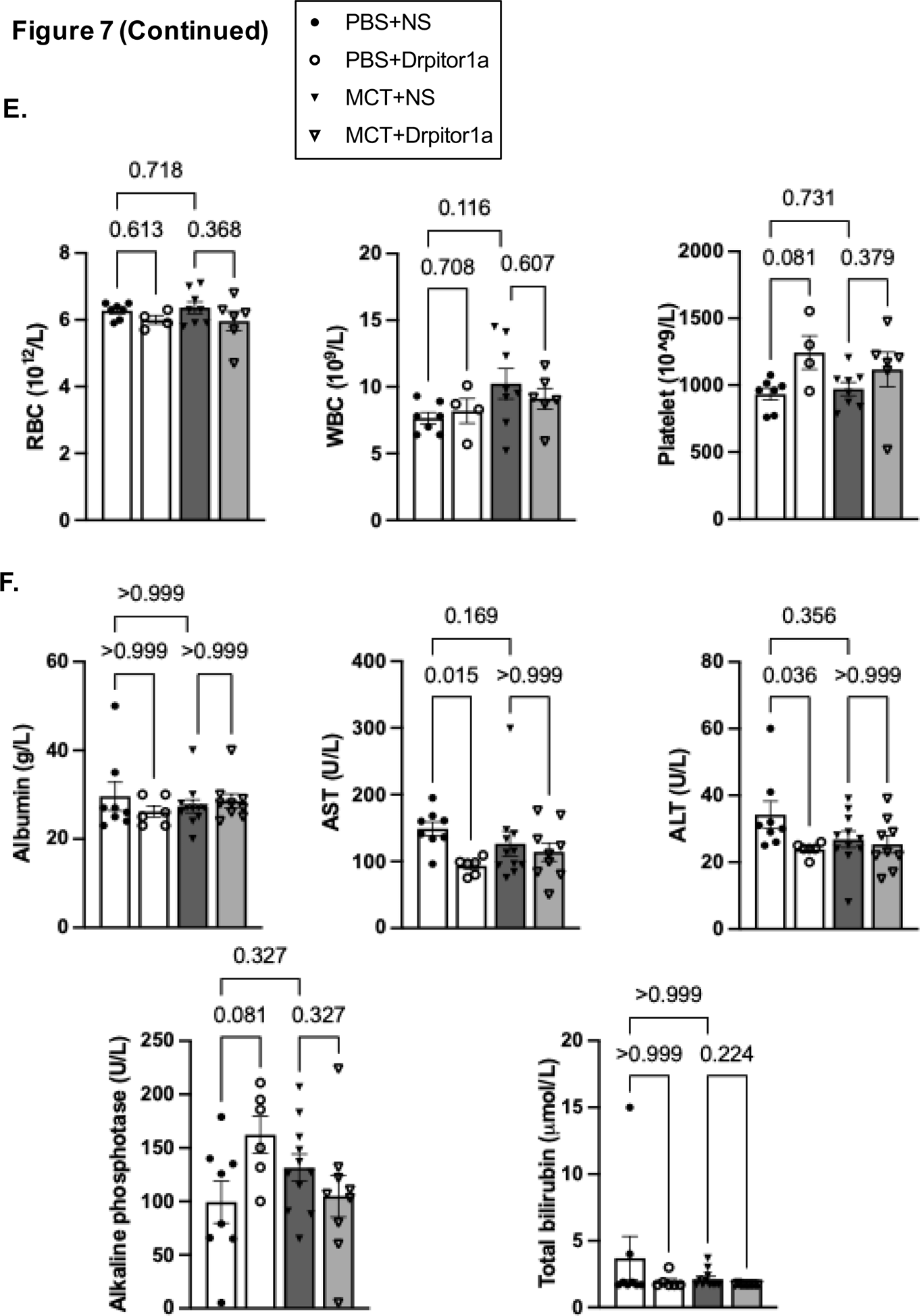
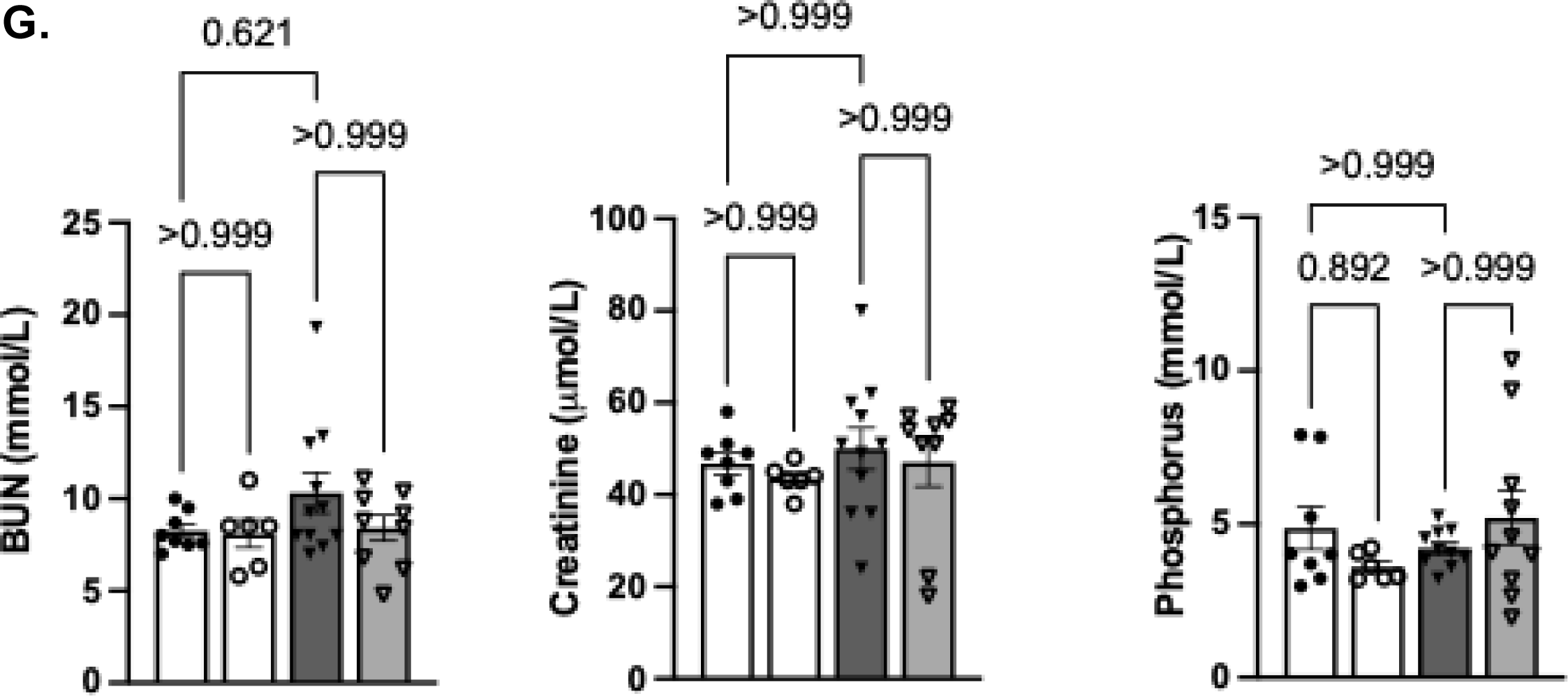
Drpitor1a regressed MCT-PAH without causing systemic toxicity in MCT-PAH rats. A. Drpitor1a treatment improved pulmonary hypertension and RV function, which is impaired in MCT-PAH rats as shown by echocardiography on day 28. MCT+NS rats had shortened PAAT, increased RVFW thickness, reduced RVFW thickening, reduced TAPSE and decreased CI compared to the PBS+NS rats. Each of these parameters was improved in the MCT+Drpitor1a group. **B. Drpitor1a treatment reduced pulmonary vascular resistance and improved RV function in MCT-PAH rats as shown by right heart catheterization on day 29.** MCT+NS rats had increased right ventricular systolic pressure (RVSP), decreased CI and increased pulmonary vascular resistance index () compared to PBS+NS rats. Drpitor1a improved CI and reduced PVR in the MCT-PAH rats (n=5-8 for PBS+NS, n=6 for PBS+Drpitor1a, n=10-11 for MCT+NS, n=9 for NS+Drpitor1a). **C. Drpitor1a treatment reduced RV hypertrophy in PAH rats induced by MCT.** RV hypertrophy was measure by the Fulton index (RV/LV+S) (n=8 for PBS+NS, n=6 for PBS+Drpitor1a, n=11 for MCT+NS, n=9 for NS+Drpitor1a). **D. Drpitor1a did not improve RV-PA coupling in MCT-PAH rats.** n=8 for PBS+NS, n=6 for PBS+Drpitor1a, n=10 for MCT+NS, n=9 for NS+Drpitor1a. **E. Drpitor1a did not induce significant hematological toxicity in PBS or MCT rats.** RBC: red blood cell; WBC: white blood cell (n=7 for PBS+NS, n=4 for PBS+Drpitor1a, n=8 for MCT+NS, n=6 for NS+Drpitor1a). **F. Drpitor1a did not induce significant hepatic toxicity in PBS or MCT rats.** AST: aspartate aminotransferase; ALT: alanine transaminase (n=8 for PBS+NS, n=6 for PBS+Drpitor1a, n=11 for MCT+NS, n=9 for NS+Drpitor1a). **G. Drpitor1a did not induce significant renal toxicity in PBS or MCT rats.** BUN: blood urea nitrogen (n=8 for PBS+NS, n=6 for PBS+Drpitor1a, n=11 for MCT+NS, n=9 for NS+Drpitor1a).

### Drpitor1a is safe for control and PAH rats

Drpitor1a did not have a detectable effect on pulmonary vascular remodeling or RV function in control rats (**Figure 7A-C**). Furthermore, Drpitor1a also did not cause significant hematological, hepatic, or renal toxicity in either control or MCT-PAH rats (**Figure 7E-G**).

## Discussion

We investigated the therapeutic efficacy of a novel, small molecule, Drp1 GTPase inhibitor in human and experimental PAH. This study has five major findings:

1. Drpitor1a inhibits the GTPase activity of recombinant and endogenous hPASMC Drp1 and prevents mitochondrial fission without blocking mitochondrial translocation of Drp1.
2. Drp1 GTPase activity is increased in PAH hPASMC and Drpitor1a reverses key hallmarks of PAH pathophysiology in PAH hPASMC, notably decreasing mitochondrial fission and reducing cell proliferation while increasing apoptosis.
3. Drpitor1a levels are significantly higher in female versus male RVs and tend to be higher in female versus male lungs.
4. Drpitor1a regresses established MCT-PAH in females, reversing adverse pulmonary vascular remodeling and, as a result, improves right heart function. There is a sexual dimorphism and Drpitor1a is most effective in females, at the dose used.
5. Drpitor1a’s beneficial effects were achieved without hepatic, renal or hematologic toxicity.

Drpitor1a is relatively selective for cells in which Drp1 GTPase is activated with no significant effect in normal hPASMC or normal rats.

PAH hPASMC have a hyperproliferative phenotype that shares several hallmarks of cancer cells(25), including increased mitochondrial fission and decreased mitochondrial fusion(26), a metabolic shift to Warburg metabolism(27), increased DNA damage(28) and resistance to apoptosis(18). Drpitor1a inhibited the Drp1 GTPase activity of human recombinant Drp1 (**Figure 1A**), demonstrating that it is a direct Drp1 GTPase inhibitor. Consistent with findings in cancer cells(22), Drpitor1a inhibited mitochondrial fission, which reversed the hyperproliferative and apoptosis-resistant disease phenotype in PAH hPASMC (**Figure 1**). Interestingly, the effective concentration of Drpitor1a for inhibiting mitochondrial fission is 5-fold lower for PAH hPASMC (0.1 μM) than for A549 cells (0.5 μM)(22). This may reflect lower intrinsic Drp1 activity in the PAH hPASMCs versus cancer cells. These findings reveal a key role for Drp1 and mitochondrial fission in PAH pathophysiology and are consistent with prior findings(19).

In cancer(20, 29, 30) and hyperproliferative PAH(19) cells, increased Drp1 expression and/or activation accelerate fission, which is permissive of rapid mitosis(9). Coordination of fission with mitosis is achieved, in part, by a reliance of both fission and cell cycle progression on shared kinases, such as cyclin B-CDK1(9). In the PAH RV, there is also Drp1 activation in RV fibroblasts, which permits rapid fibroblast proliferation and increased collagen production(31). In RV cardiomyocytes, fission triggered by RV ischemia and reperfusion increases ROS production and impairs RV function(32).

*In vivo*, mdivi-1 regresses pulmonary hypertension in a rat model of PAH, induced by exposure to the HIF-1α activator, cobalt chloride(19). While mdivi-1 is beneficial for the pulmonary vasculature and RV in PAH, concerns have been raised regarding its specificity for Drp1. Reported off-target effects of mdivi-1 include changes in the function of the mitochondrial electron transport chain complex I(33, 34) and L-OPA1(34). Whether these effects of mdivi-1 are truly off-target or related to Drp1 inhibition is unclear; but the controversy confounds interpretation of studies using mdivi-1. This led us to develop new, potent and specific Drp1 GTPase inhibitors(22), Drpitor and Drpitor1a (the latter modified by the removal of a methoxymethyl group to reduce hydrolytic lability)(22).

The half-life (IV) and oral bioavailability of Drpitor1a is lower in females than males (**Table 1**). However, at the dose used in the regression study (1 mg/kg), females have higher concentrations of Drpitor1a in RV tissues, and tend to have higher lung tissue levels, irrespective of the route of administration (**Figure 3**). Due to the low oral bioavailability of Drpitor1a (1mg/kg) (female 12.6% versus male 26.0%), we administered Drpitor1a IV (1mg/kg) for *in vivo* studies.

In the pilot study, males developed more severe pulmonary vascular remodeling and RV hypertrophy than females (**Figure 4**), as reported(35, 36). This sex difference may be explained by the fact that MCT is a pyrrolizidine alkaloid, the toxicity of which is determined by cytochrome P450 (CYP450)-induced bioactivation in the liver (37). In rats, CYP3A is the primary mediator of MCT oxidation, producing dehydro-MCT(37). Male rat livers generate higher basal bioactivation of MCT than do female livers(38). The sexual dimorphism in MCT metabolism merits further investigation as a potential explanation for the more severe PAH in males in this model.

In the pilot study, Drpitor1a showed a trend to improve pulmonary vascular remodeling and RV hypertrophy only in female MCT-PAH rats (**Figure 4**). This greater therapeutic efficacy in females may be explained by the higher effective tissue concentrations of Drpitor1a (**Figure 2-3**). Sex differences in drug pharmacokinetics, which are well described(39–41), may relate to differences in drug absorption, distribution, metabolism, and elimination(42). Differences in drug metabolism commonly contribute to sex differences in drug pharmacokinetics(43). Various CYP450 superfamily members have sex-specific activity levels(44), although these differences vary by enzyme and drug (44–46). Fluctuation of sex hormones in females during the menstrual cycle alter the activity of CYP450 enzymes(47). While we identify a sexual dimorphism in the response to Drpitor1a, further research is required to determine the basis for the observed pharmacokinetic differences and to establish whether sex differences could be overcome by sex-specific dosing.

Day 14 post-MCT echocardiography confirmed that pulmonary hypertension and RV dysfunction had developed (**Figure 5B-F**). Therefore, the Drpitor1a administered from days 17 to 27 post- MCT can be viewed as a regression therapy, making it more relevant to the clinical scenario. PAH worsened significantly between days 14 and 28 in the MCT+NS group, evident as a significant decrease in PAAT, while the RV deteriorated, indicated by increases in RVFW thickness and decreases in RVFW thickening, TAPSE, and CI (**Figure E7**). However, in the MCT+Drpitor1a group, there was no significant difference in these parameters over this period, indicating that Drpitor1a reduces PH and preserves RV function in MCT-PAH.

We considered whether Drpitor1a’s benefits relate to regression of pulmonary vascular disease versus an independent improvement in RV function. RV-PA coupling defined as the ratio of end- systolic to arterial elastances (Ees/Ea)(48) is an accepted measurement of RV adaptation to afterload. In patients with PH, decreased RV-PA coupling predicts RV failure(49). We used validated surrogates for Ees/Ea (TAPSE/mPAP and RVFW thickening/mPAP) in this study(24) but found no significant increase in either parameter in the MCT+Drpitor1a group (**Figure 7D**).

Thus, Drpitor1a primarily improves pulmonary vascular remodeling and secondarily improves RV function by reducing RV afterload, which is in contrast with several mitochondrial metabolic therapies for PAH. For example, Tian et al. discovered that targeting aerobic glycolysis in RV fibroblasts using dichloroacetate, a pyruvate dehydrogenase kinase inhibitor, can effectively improve RV-PA coupling in MCT-PAH rats(24). Dichloroacetate also regresses pulmonary vascular disease in some patients with PAH(50).

### Study limitations

Due to the concordant, positive, effects of Drpitor1a in PAH hPASMC and the MCT model, as well as the cost of Drpitor1a, which we synthesized ourselves, we did not explore Drpitor1a in the SU5416-hypoxia PAH model. We did not assess the therapeutic efficacy of Drpitor1a in male rats at a dose higher than 1mg/kg and thus cannot exclude the possibility that Drpitor1a may have therapeutic efficacy in male rats at higher doses.

## Conclusions

GTPase activity is increased in PAH hPASMC. Inhibition of Drp1 by Drpitor1a prevents mitochondrial fission, reduces PAH hPASMC proliferation *ex vivo* and regresses MCT- PAH *in vivo.* Drpitor1a had no significant effects on normal hPASMC and *in vivo* caused no toxicity at the dose and duration used. There is a sexual dimorphism in the pharmacokinetics and therapeutic efficacy of this novel Drp1 inhibitor which merits further exploration.

## Acknowledgments

This research is supported by the Translational Institute of Medicine (TIME) and Queen’s Cardiopulmonary Unit (QCPU) at Queen’s University.

## Author contributions

D. Wu, A. Dasgupta, and S. L. Archer designed the research; R. D. Jansen-van Vuuren and L. Jedlovčnik synthesized Drpitor1a; D. Wu, A. Dasgupta, R. Al-Qazazi, K-H Chen, A. Martin, J. D. Mewburn, E. Alizadeh, P. D.A. Lima, O. Jones, P. Colpman, N. Breault, and I. Emon performed the research; D. Wu and P. D.A. Lima analyzed the data; Y.Y. Zhao and G. Sutendra analyzed the tissue concentration of Drpitor1a; D. Wu, A. Dasgupta, and S. L. Archer wrote the manuscript; S. L. Archer supervised the experiments and acquired funding.

## Conflict of interest

The authors acknowledge that they hold a patent for the use of Drpitor1a in diseases including pulmonary hypertension and cancer: Wu D, Wells M, Archer SL. Inhibitors of Mitochondrial Fission. Patent Application No. 62/826,247.

Non-standard Abbreviations
|  |  |
| --- | --- |
| CI | Cardiac index |
| CO | Cardiac output |
| Drpitor | Drp1 inhibitor |
| Drp1 | Dynamin-related protein 1 |
| ER | Endoplasmic reticulum |
| HR | Heart rate |
| ID | Inner diameter |
| LJV | Left jugular vein |
| LVEDP | Left ventricular end-diastolic pressure |
| MCT | Monocrotaline |
| MFC | Mitochondrial fragmentation count |
| mPAP | Mean pulmonary artery pressure |
| NS | Normal saline |
| OMM | Outer mitochondrial membrane |
| PAAT | Pulmonary artery acceleration time |
| PA | Pulmonary artery |
| PAH | Pulmonary arterial hypertension |
| PASMC | Pulmonary artery smooth muscle cell |
| PVR | Pulmonary vascular resistance |
| RV | Right ventricle |
| RVFW | Right ventricular free wall |
| RVSP | Right ventricular systolic pressure |
| RHC | Right heart catheterization |
| TAPSE | Tricuspid annular plane systolic excursion |
| TMRM | Tetramethylrhodamine methyl ester |
| VTI | Velocity time integral |

## References

1. Mair KM, Johansen AK, Wright AF, Wallace E, MacLean MR. Pulmonary arterial hypertension: basis of sex differences in incidence and treatment response. Br J Pharmacol 2014; 171: 567–579.

2. Humbert M, Guignabert C, Bonnet S, Dorfmuller P, Klinger JR, Nicolls MR, Olschewski AJ, Pullamsetti SS, Schermuly RT, Stenmark KR, Rabinovitch M. Pathology and pathobiology of pulmonary hypertension: state of the art and research perspectives. Eur Respir J 2019; 53.

3. Simonneau G, Montani D, Celermajer DS, Denton CP, Gatzoulis MA, Krowka M, Williams PG, Souza R. Haemodynamic definitions and updated clinical classification of pulmonary hypertension. Eur Respir J 2019; 53.

4. Al-Qazazi R, Lima PDA, Prisco SZ, Potus F, Dasgupta A, Chen KH, Tian L, Bentley RET, Mewburn J, Martin AY, Wu D, Jones O, Maurice DH, Bonnet S, Provencher S, Prins KW, Archer SL. Macrophage-NLRP3 Activation Promotes Right Ventricle Failure in Pulmonary Arterial Hypertension. Am J Respir Crit Care Med 2022; 206: 608–624.

5. Ryan JJ, Archer SL. The right ventricle in pulmonary arterial hypertension: disorders of metabolism, angiogenesis and adrenergic signaling in right ventricular failure. Circ Res 2014; 115: 176–188.

6. Prins KW, Thenappan T, Weir EK, Kalra R, Pritzker M, Archer SL. Repurposing Medications for Treatment of Pulmonary Arterial Hypertension: What’s Old Is New Again. J Am Heart Assoc 2019; 8: e011343.

7. Thenappan T, Ormiston ML, Ryan JJ, Archer SL. Pulmonary arterial hypertension: pathogenesis and clinical management. BMJ 2018; 360: j5492.

8. Thenappan T, Ryan JJ, Archer SL. Evolving epidemiology of pulmonary arterial hypertension. Am J Respir Crit Care Med 2012; 186: 707–709.

9. Archer SL. Mitochondrial dynamics--mitochondrial fission and fusion in human diseases. N Engl J Med 2013; 369: 2236–2251.

10. Breault NM, Wu D, Dasgupta A, Chen KH, Archer SL. Acquired disorders of mitochondrial metabolism and dynamics in pulmonary arterial hypertension. Front Cell Dev Biol 2023; 11: 1105565.

11. Archer SL, Marsboom G, Kim GH, Zhang HJ, Toth PT, Svensson EC, Dyck JR, Gomberg-Maitland M, Thebaud B, Husain AN, Cipriani N, Rehman J. Epigenetic attenuation of mitochondrial superoxide dismutase 2 in pulmonary arterial hypertension: a basis for excessive cell proliferation and a new therapeutic target. Circulation 2010; 121: 2661–2671.

12. Caruso P, Dunmore BJ, Schlosser K, Schoors S, Dos Santos C, Perez-Iratxeta C, Lavoie JR, Zhang H, Long L, Flockton AR, Frid MG, Upton PD, D’Alessandro A, Hadinnapola C, Kiskin FN, Taha M, Hurst LA, Ormiston ML, Hata A, Stenmark KR, Carmeliet P, Stewart DJ, Morrell NW. Identification of MicroRNA-124 as a Major Regulator of Enhanced Endothelial Cell Glycolysis in Pulmonary Arterial Hypertension via PTBP1 (Polypyrimidine Tract Binding Protein) and Pyruvate Kinase M2. Circulation 2017; 136: 2451–2467.

13. Zhang H, Wang D, Li M, Plecita-Hlavata L, D’Alessandro A, Tauber J, Riddle S, Kumar S, Flockton A, McKeon BA, Frid MG, Reisz JA, Caruso P, El Kasmi KC, Jezek P, Morrell NW, Hu CJ, Stenmark KR. Metabolic and Proliferative State of Vascular Adventitial Fibroblasts in Pulmonary Hypertension Is Regulated Through a MicroRNA-124/PTBP1 (Polypyrimidine Tract Binding Protein 1)/Pyruvate Kinase Muscle Axis. Circulation 2017; 136: 2468–2485.

14. Sutendra G, Michelakis ED. The metabolic basis of pulmonary arterial hypertension. Cell Metab 2014; 19: 558-573.

15. Smirnova E, Griparic L, Shurland DL, van der Bliek AM. Dynamin-related protein Drp1 is required for mitochondrial division in mammalian cells. Mol Biol Cell 2001; 12: 2245–2256.

16. Loson OC, Song Z, Chen H, Chan DC. Fis1, Mff, MiD49, and MiD51 mediate Drp1 recruitment in mitochondrial fission. Mol Biol Cell 2013; 24: 659-667.

17. Yoon Y, Pitts KR, McNiven MA. Mammalian dynamin-like protein DLP1 tubulates membranes. Mol Biol Cell 2001; 12: 2894–2905.

18. Chen KH, Dasgupta A, Lin J, Potus F, Bonnet S, Iremonger J, Fu J, Mewburn J, Wu D, Dunham- Snary K, Theilmann AL, Jing ZC, Hindmarch C, Ormiston ML, Lawrie A, Archer SL. Epigenetic Dysregulation of the Dynamin-Related Protein 1 Binding Partners MiD49 and MiD51 Increases Mitotic Mitochondrial Fission and Promotes Pulmonary Arterial Hypertension: Mechanistic and Therapeutic Implications. Circulation 2018; 138: 287–304.

19. Marsboom G, Toth PT, Ryan JJ, Hong Z, Wu X, Fang YH, Thenappan T, Piao L, Zhang HJ, Pogoriler J, Chen Y, Morrow E, Weir EK, Rehman J, Archer SL. Dynamin-related protein 1- mediated mitochondrial mitotic fission permits hyperproliferation of vascular smooth muscle cells and offers a novel therapeutic target in pulmonary hypertension. Circ Res 2012; 110: 1484–1497.

20. Rehman J, Zhang HJ, Toth PT, Zhang Y, Marsboom G, Hong Z, Salgia R, Husain AN, Wietholt C, Archer SL. Inhibition of mitochondrial fission prevents cell cycle progression in lung cancer. FASEB J 2012; 26: 2175–2186.

21. Cassidy-Stone A, Chipuk JE, Ingerman E, Song C, Yoo C, Kuwana T, Kurth MJ, Shaw JT, Hinshaw JE, Green DR, Nunnari J. Chemical inhibition of the mitochondrial division dynamin reveals its role in Bax/Bak-dependent mitochondrial outer membrane permeabilization. Dev Cell 2008; 14: 193–204.

22. Wu D, Dasgupta A, Chen KH, Neuber-Hess M, Patel J, Hurst TE, Mewburn JD, Lima PDA, Alizadeh E, Martin A, Wells M, Snieckus V, Archer SL. Identification of novel dynamin- related protein 1 (Drp1) GTPase inhibitors: Therapeutic potential of Drpitor1 and Drpitor1a in cancer and cardiac ischemia-reperfusion injury. FASEB J 2020; 34: 1447–1464.

23. Hong Z, Chen KH, DasGupta A, Potus F, Dunham-Snary K, Bonnet S, Tian L, Fu J, Breuils-Bonnet S, Provencher S, Wu D, Mewburn J, Ormiston ML, Archer SL. MicroRNA-138 and MicroRNA-25 Down-regulate Mitochondrial Calcium Uniporter, Causing the Pulmonary Arterial Hypertension Cancer Phenotype. Am J Respir Crit Care Med 2017; 195: 515–529.

24. Tian L, Wu D, Dasgupta A, Chen KH, Mewburn J, Potus F, Lima PDA, Hong Z, Zhao YY, Hindmarch CCT, Kutty S, Provencher S, Bonnet S, Sutendra G, Archer SL. Epigenetic Metabolic Reprogramming of Right Ventricular Fibroblasts in Pulmonary Arterial Hypertension: A Pyruvate Dehydrogenase Kinase-Dependent Shift in Mitochondrial Metabolism Promotes Right Ventricular Fibrosis. Circ Res 2020; 126: 1723–1745.

25. Boucherat O, Vitry G, Trinh I, Paulin R, Provencher S, Bonnet S. The cancer theory of pulmonary arterial hypertension. Pulm Circ 2017; 7: 285–299.

26. Chen H, Chan DC. Mitochondrial Dynamics in Regulating the Unique Phenotypes of Cancer and Stem Cells. Cell Metab 2017; 26: 39–48.

27. Archer SL. Pyruvate Kinase and Warburg Metabolism in Pulmonary Arterial Hypertension: Uncoupled Glycolysis and the Cancer-Like Phenotype of Pulmonary Arterial Hypertension. Circulation 2017; 136: 2486–2490.

28. Meloche J, Pflieger A, Vaillancourt M, Paulin R, Potus F, Zervopoulos S, Graydon C, Courboulin A, Breuils-Bonnet S, Tremblay E, Couture C, Michelakis ED, Provencher S, Bonnet S. Role for DNA damage signaling in pulmonary arterial hypertension. Circulation 2014; 129: 786–797.

29. Zhao J, Zhang J, Yu M, Xie Y, Huang Y, Wolff DW, Abel PW, Tu Y. Mitochondrial dynamics regulates migration and invasion of breast cancer cells. Oncogene 2013; 32: 4814–4824.

30. Xie Q, Wu Q, Horbinski CM, Flavahan WA, Yang K, Zhou W, Dombrowski SM, Huang Z, Fang X, Shi Y, Ferguson AN, Kashatus DF, Bao S, Rich JN. Mitochondrial control by DRP1 in brain tumor initiating cells. Nat Neurosci 2015; 18: 501–510.

31. Tian L, Potus F, Wu D, Dasgupta A, Chen KH, Mewburn J, Lima P, Archer SL. Increased Drp1- Mediated Mitochondrial Fission Promotes Proliferation and Collagen Production by Right Ventricular Fibroblasts in Experimental Pulmonary Arterial Hypertension. Front Physiol 2018; 9: 828.

32. Tian L, Neuber-Hess M, Mewburn J, Dasgupta A, Dunham-Snary K, Wu D, Chen KH, Hong Z, Sharp WW, Kutty S, Archer SL. Ischemia-induced Drp1 and Fis1-mediated mitochondrial fission and right ventricular dysfunction in pulmonary hypertension. J Mol Med (Berl) 2017; 95: 381–393.

33. Bordt EA, Clerc P, Roelofs BA, Saladino AJ, Tretter L, Adam-Vizi V, Cherok E, Khalil A, Yadava N, Ge SX, Francis TC, Kennedy NW, Picton LK, Kumar T, Uppuluri S, Miller AM, Itoh K, Karbowski M, Sesaki H, Hill RB, Polster BM. The Putative Drp1 Inhibitor mdivi-1 Is a Reversible Mitochondrial Complex I Inhibitor that Modulates Reactive Oxygen Species. Dev Cell 2017; 40: 583–594 e586.

34. Aishwarya R, Alam S, Abdullah CS, Morshed M, Nitu SS, Panchatcharam M, Miriyala S, Kevil CG, Bhuiyan MS. Pleiotropic effects of mdivi-1 in altering mitochondrial dynamics, respiration, and autophagy in cardiomyocytes. Redox Biol 2020; 36: 101660.

35. Vijay P, Tighe SM, Jha D, Sharp TG, Brown JW. P-103: Gender differences in the development of pulmonary hypertension. American Journal of Hypertension 2001; 14: 64A–64A.

36. Bal E, Ilgin S, Atli O, Ergun B, Sirmagul B. The effects of gender difference on monocrotaline-induced pulmonary hypertension in rats. Hum Exp Toxicol 2013; 32: 766–774.

37. Fu PP, Xia Q, Lin G, Chou MW. Pyrrolizidine alkaloids--genotoxicity, metabolism enzymes, metabolic activation, and mechanisms. Drug Metab Rev 2004; 36: 1–55.

38. Luo J, Yang X, Qiu S, Li X, Xiang E, Fang Y, Wang Y, Zhang L, Wang H, Zheng J, Guo Y. Sex difference in monocrotaline-induced developmental toxicity and fetal hepatotoxicity in rats. Toxicology 2019; 418: 32–40.

39. Zucker I, Prendergast BJ. Sex differences in pharmacokinetics predict adverse drug reactions in women. Biol Sex Differ 2020; 11: 32.

40. Fadiran EO, Zhang L. Effects of Sex Differences in the Pharmacokinetics of Drugs and Their Impact on the Safety of Medicines in Women. In: Harrison-Woolrych M, editor. Medicines For Women. Cham: Springer International Publishing; 2015. p. 41-68.

41. Kokras N, Dalla C, Papadopoulou-Daifoti Z. Sex differences in pharmacokinetics of antidepressants. Expert Opin Drug Metab Toxicol 2011; 7: 213–226.

42. Gandhi M, Aweeka F, Greenblatt RM, Blaschke TF. Sex differences in pharmacokinetics and pharmacodynamics. Annu Rev Pharmacol Toxicol 2004; 44: 499–523.

43. Waxman DJ, Holloway MG. Sex differences in the expression of hepatic drug metabolizing enzymes. Mol Pharmacol 2009; 76: 215–228.

44. Scandlyn MJ, Stuart EC, Rosengren RJ. Sex-specific differences in CYP450 isoforms in humans. Expert Opin Drug Metab Toxicol 2008; 4: 413–424.

45. Higashi E, Fukami T, Itoh M, Kyo S, Inoue M, Yokoi T, Nakajima M. Human CYP2A6 is induced by estrogen via estrogen receptor. Drug Metab Dispos 2007; 35: 1935–1941.

46. Yang L, Li Y, Hong H, Chang CW, Guo LW, Lyn-Cook B, Shi L, Ning B. Sex Differences in the Expression of Drug-Metabolizing and Transporter Genes in Human Liver. J Drug Metab Toxicol 2012; 3: 1000119.

47. Asprodini E, Tsiokou V, Begas E, Kilindris T, Kouvaras E, Samara M, Messinis I. Alterations in Xenobiotic-Metabolizing Enzyme Activities across Menstrual Cycle in Healthy Volunteers. J Pharmacol Exp Ther 2019; 368: 262–271.

48. Sunagawa K, Maughan WL, Burkhoff D, Sagawa K. Left ventricular interaction with arterial load studied in isolated canine ventricle. Am J Physiol 1983; 245: H773–780.

49. Tello K, Dalmer A, Axmann J, Vanderpool R, Ghofrani HA, Naeije R, Roller F, Seeger W, Sommer N, Wilhelm J, Gall H, Richter MJ. Reserve of Right Ventricular-Arterial Coupling in the Setting of Chronic Overload. Circ Heart Fail 2019; 12: e005512.

50. Michelakis ED, Gurtu V, Webster L, Barnes G, Watson G, Howard L, Cupitt J, Paterson I, Thompson RB, Chow K, O’Regan DP, Zhao L, Wharton J, Kiely DG, Kinnaird A, Boukouris AE, White C, Nagendran J, Freed DH, Wort SJ, Gibbs JSR, Wilkins MR. Inhibition of pyruvate dehydrogenase kinase improves pulmonary arterial hypertension in genetically susceptible patients. Sci Transl Med 2017; 9.

