## Supplementary material for "Efficacy of Drpitor1a, a Dynamin-Related Protein 1 inhibitor, in Pulmonary Arterial Hypertension": Online Data Supplement

**Figure E1**

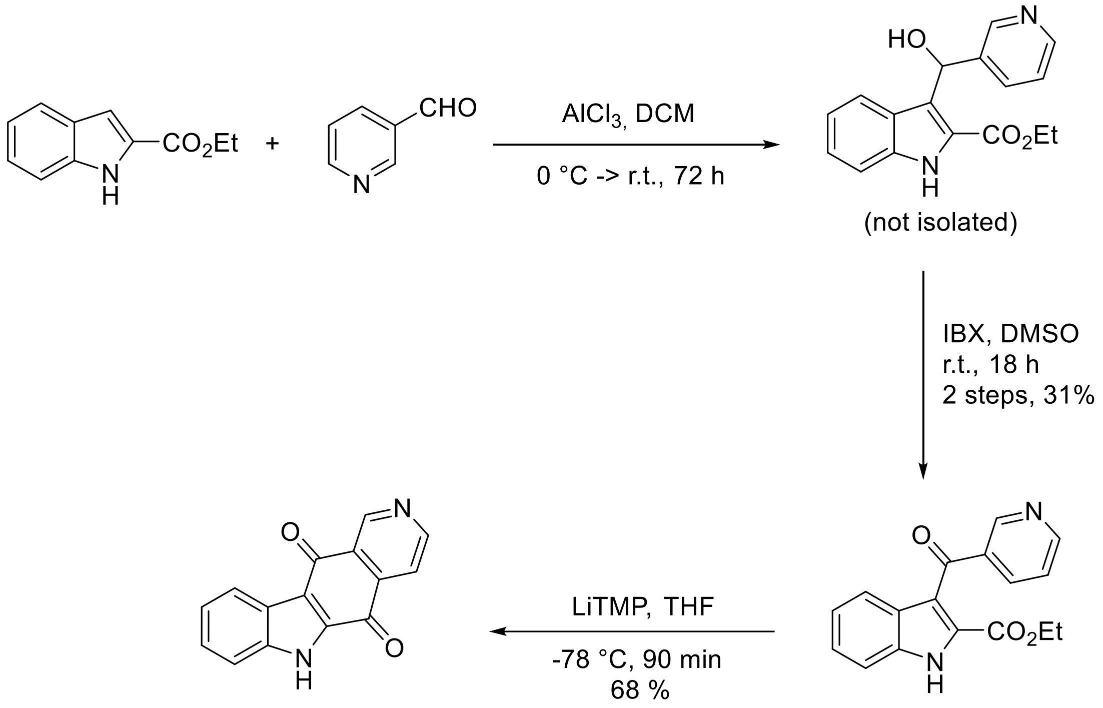


**Figure E1.** A modified reaction scheme of Drpitor1a synthesis. The procedure was modified from Ramkumar&Nagarajan, DOI: 10.1021/jo402593w

**Figure E2**

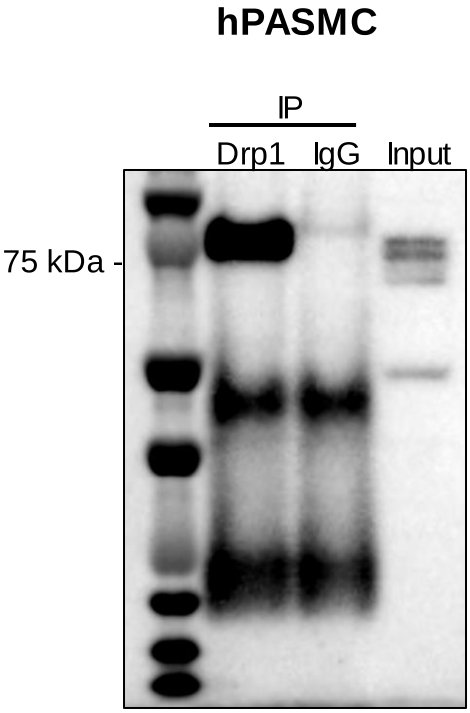


**Figure E2.** Immunoblot showing Drp1 protein was specifically immunoprecipited by Drp1 antibody in human PASMC.

#### Figure E3

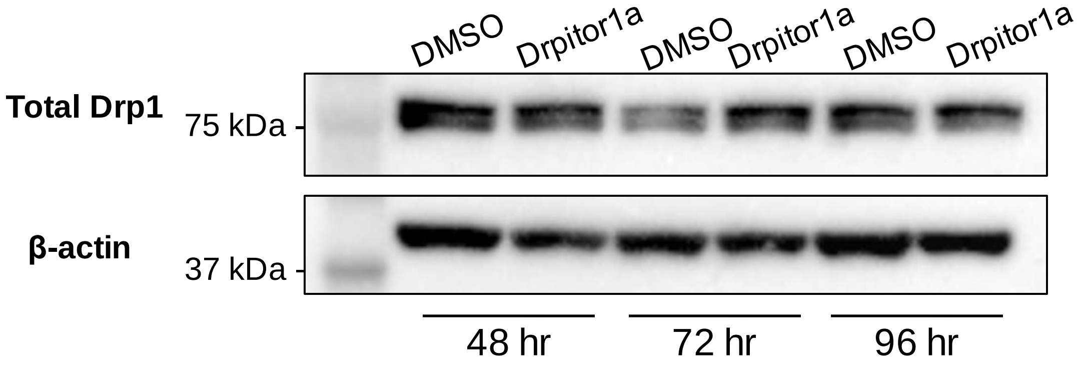


**Figure E3.** Immunoblot showing Drpitor1a does not reduce global Drp1 expression in PAH hPASMC.Representative immunoblot shows treatment of PASMC with Drpitor1a increases total Drp1 at 72 hours.

**Figure E4**

Bak

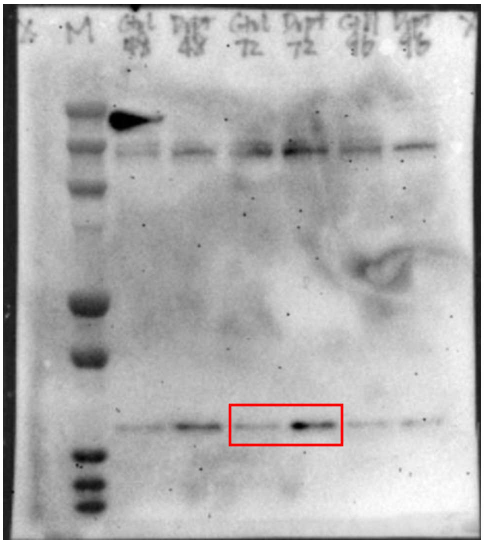

β-actin

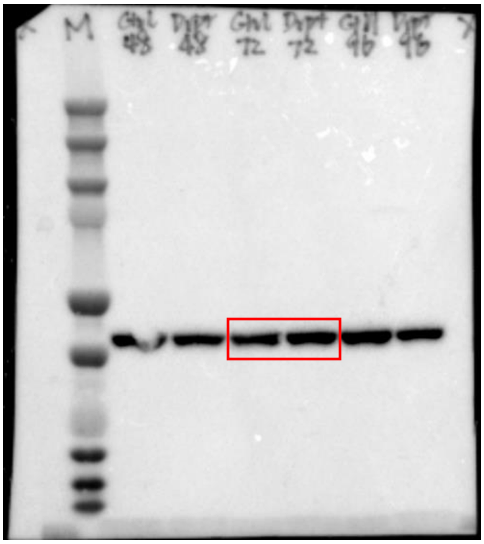

Bcl-2

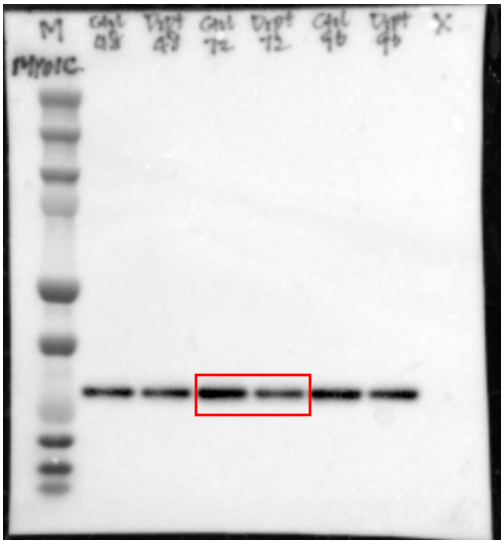

β-actin

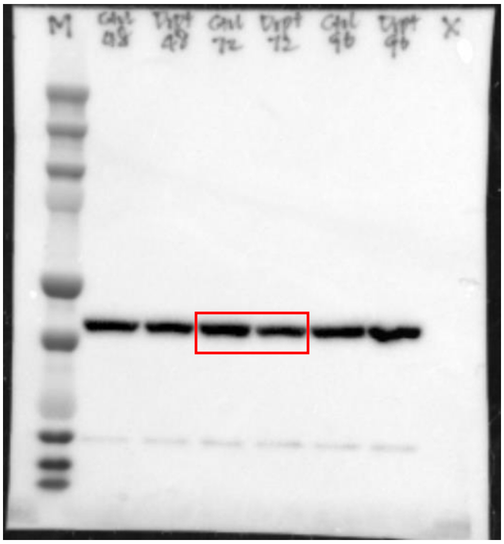

#### Figure E4 (continued)

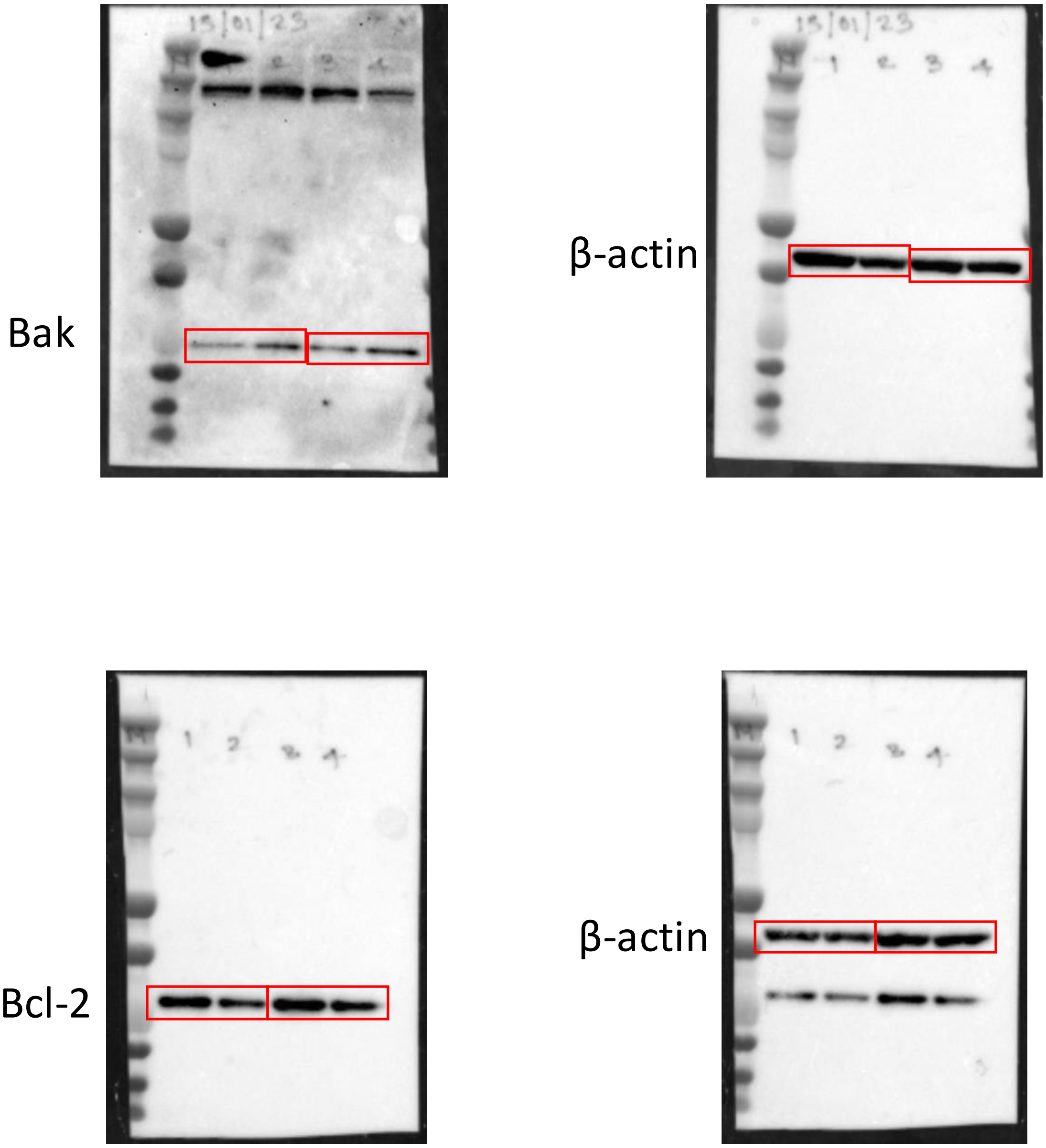


**Figure E4.** Original immunoblots for Figure 1J.

### Figure E5

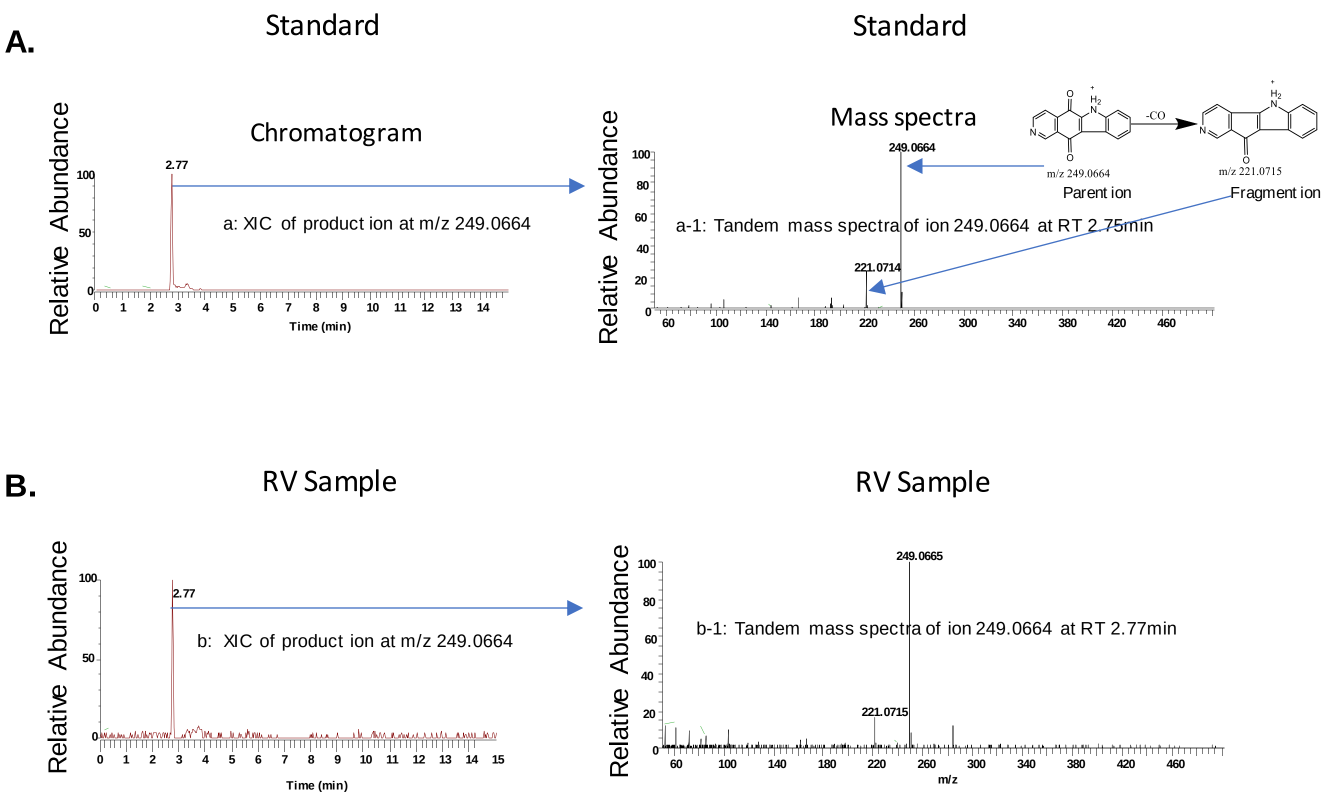


**Figure E5.** Identification of Drpitor1a by HPLC coupled with high resolution orbitrap mass spec. Extracted ion chromatogram(XIC) of ion at m/z 249.0664 from 249.0664 from product ion scan for ion at m/z 249.0664 (A) standard (B) RV sample; Tandem mass spectra of ion at m/z 249.0664 for peak at retention time 2.77min (a-1) standard, (b-1) RV sample.

Drpitor1a in RV sample was identified by HPLC coupled with high resolution Orbitrap mass spectrometry. By running standard Drptor1a and RV sample, we observed the retention time of XIC at m/z 249.0665 in sample RV PK13 from product ion scan of ion at m/z 249.0665 matches the standard (a, standard and b, RV sample). Furthermore, the tandem mass spectra of parent ion at m/z 249.0665 gave rise to a fragment ion at m/z 221.0715 with neutral loss of CO (a-1 standard and b-1 RV sample) , which shows the fingerprint of the structure. The observed parent and fragment ions from standard and RV sample also matched their theoretical value at 249.0664 and 221.0715, respectively. The retention time from HPLC, high mass accuracy obtained and the fragment ion pattern confidently confirmed the existence of Drptor1a in treated RV sample.

#### Figure E6

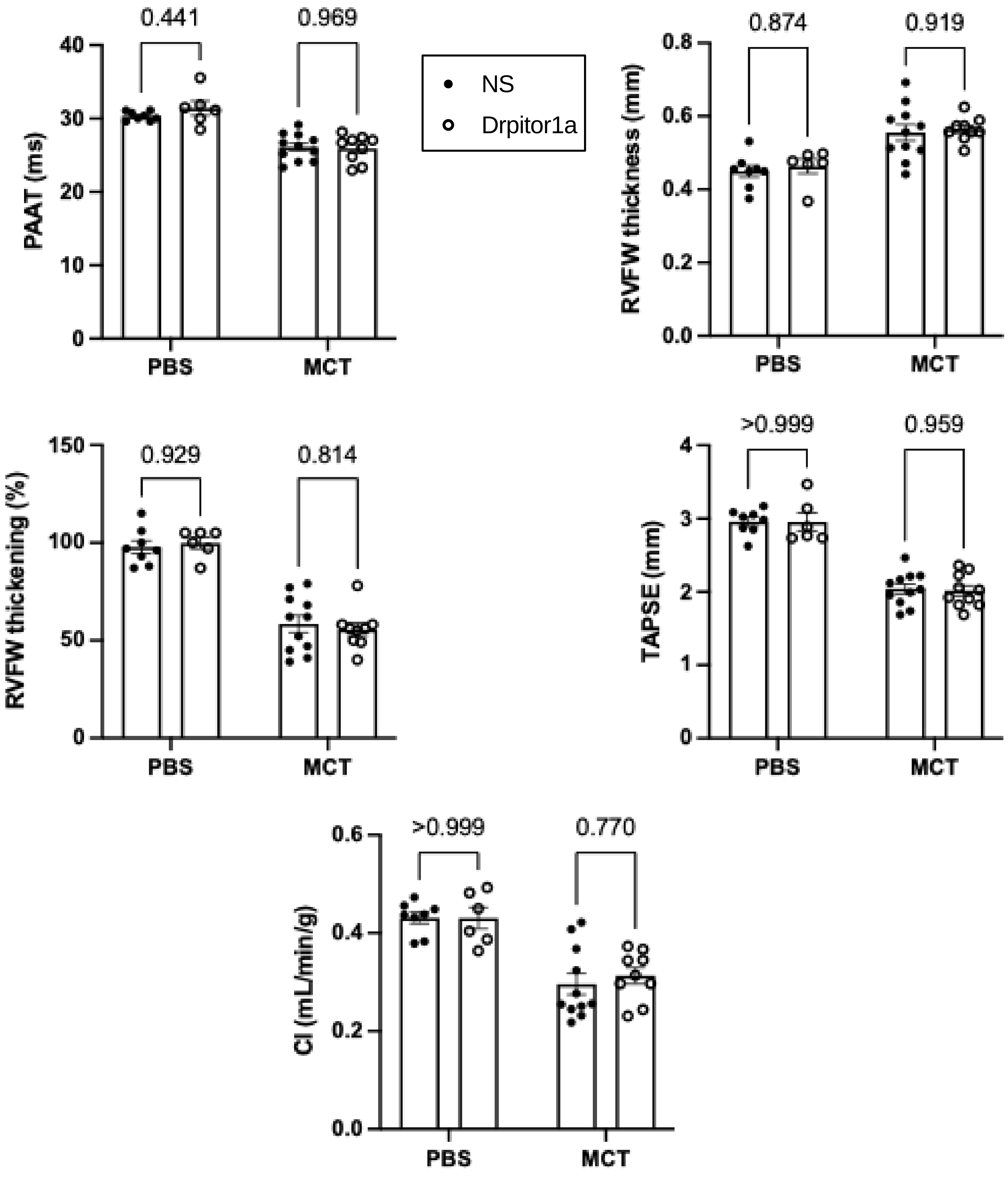


**Figure E6.** At randomization, there was no significant difference in echocardiography parameters between the rats which were randomly assigned to receive normal saline (NS) or Drpitor1a.

**Figure E7**

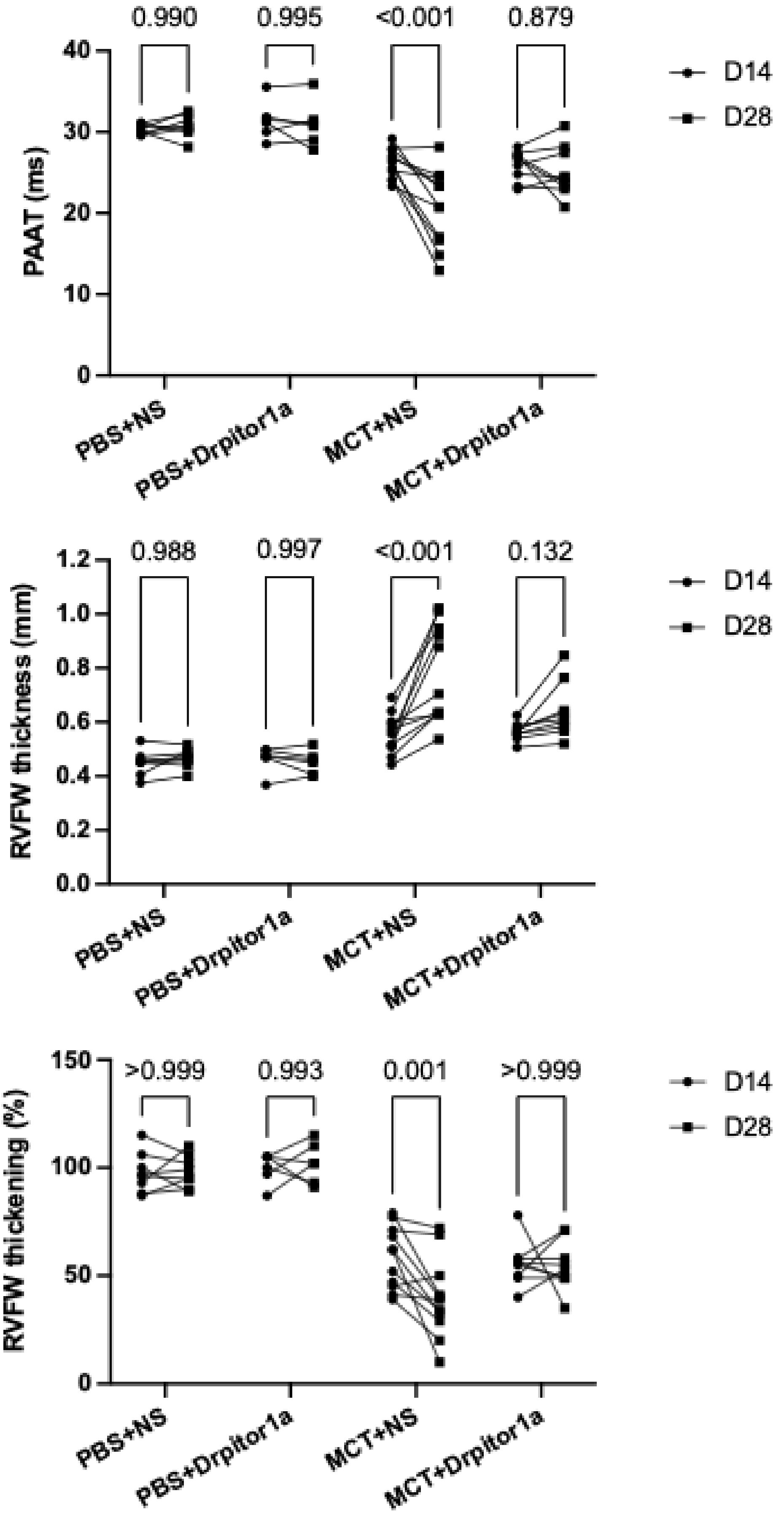
0.990
0.995
<0.001
0.879
PAAT (ms)
40
30
20
10
0
D14
D28
PBS+NS
PBS+Drpitor1a
MCT+NS
MCT+Drpitor1a
0.988
0.997
<0.001
0.132
RVFW thickness (mm)
1.2
1.0
0.8
0.6
0.4
0.2
0.0
D14
D28
PBS+NS
PBS+Drpitor1a
MCT+NS
MCT+Drpitor1a
>0.999
0.993
0.001
>0.999
RVFW thickening (%)
150
100
50
0
D14
D28
PBS+NS
PBS+Drpitor1a
MCT+NS
MCT+Drpitor1a

**Figure E7 (Continued)**

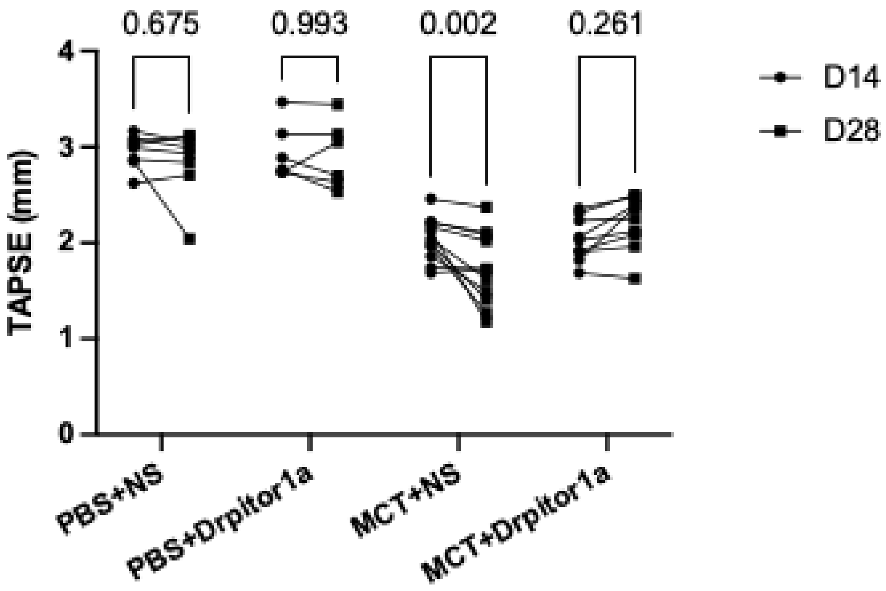


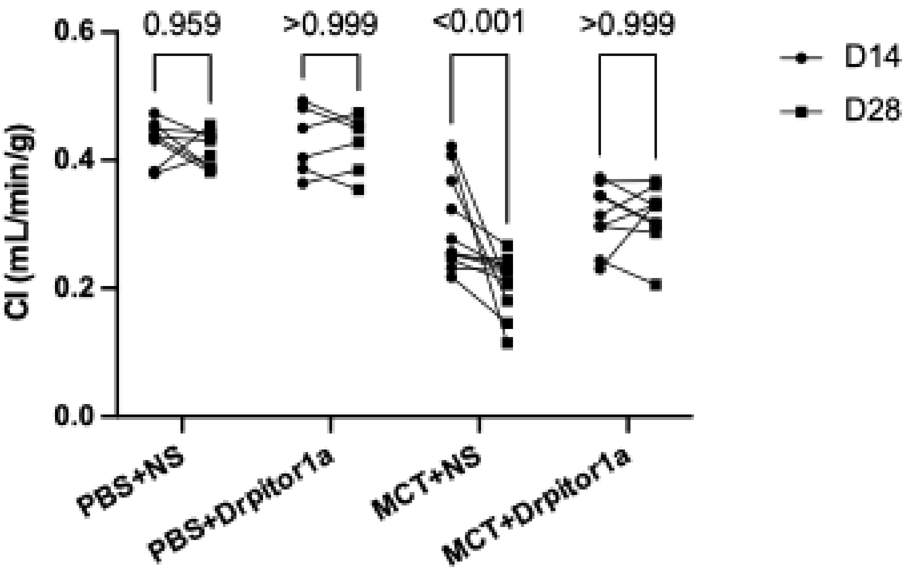


**Figure E7.** Drpitor1a preserved PAAT, RVFW thickness, RVFW thickening, TAPSE and CI in MCT-PAH rats.
