## Supplementary material for "Efficacy of Drpitor1a, a Dynamin-Related Protein 1 inhibitor, in Pulmonary Arterial Hypertension": Figure E1-E7

**Online Data Supplement**

### 1. *In vitro* studies

#### 1.1. Compound synthesis

See Figure E1.

#### 1.2. Cell Culture

Human pulmonary artery smooth muscle cells (hPASMC) from normal donors (n=3) and patients with PAH (n=5) were maintained in Human Vascular Smooth Muscle Cell Basal Medium (M231500, ThermoFisher Scientific, Burlington, ON, Canada) supplemented with Smooth Muscle Growth Supplement (S00725, ThermoFisher Scientific, Burlington, ON, Canada) and 10% heat-inactivated fetal bovine serum (12484-028, Gibco, Burlington, ON, Canada).

#### 1.3. GTPase assay

The GTPase assay is a colorimetric assay that detects phosphate. GTPase activity from both the recombinant Drp1 and immunoprecipitated Drp1 derived from PAH hPASMC was quantified using a GTPase assay kit (ab270553, Abcam, Toronto, ON, Canada) according to the manufacturer's specified protocol. OD620 was measured by a SpectraMax M3 microplate reader (Molecular Devices, LLC., San Jose, CA, USA).

In the recombinant Drp1 experiments, human recombinant Drp1 (0.5 μg in 100μL, TP321708, OriGene Technologies, Inc., Rockville, MD, USA) was first incubated with Drpitor1a (1 μM) at 4 °C for 1 hour. Then 100 μL of GTP buffer (1mM GTP in 0.1M Tris-HCl and 5mM $MgCl_2$) was added and incubated at 30℃ for 30 minutes.

In studies of Drp1 derived from PAH hPASMC, the protein was extracted using cell lysis buffer (9803, Cell Signaling Technology, Danvers, MA, USA) followed by sonication. A total of 1,000 μg of protein from each sample was allowed to bind to 2 μg of total Drp1 antibody (611112, BD Transduction Laboratories, San Jose, CA, USA) at 4℃ for 5 hours. This was followed by immunoprecipitation with 50 μL of protein A/G-agarose beads (sc-2003, Santa Cruz Biotechnology, Dallas, TX, USA) at 4℃ overnight. Then the protein-antibody-beads complex was washed twice with RIPA lysis and extraction buffer (89901, ThermoFisher Scientific, Burlington, ON, Canada), twice with GTPase buffer (50 μM Tris-HCl pH 7.5, 2.5 μM $MgCl_2$), and finally incubated with 0.5 mM GTP at 30℃ for 30 minutes.

##### 1.3. Mitochondrial morphology analysis

Cells were treated with Drpitor1 (0.1 μM) or DMSO for 4-5 hours and mitochondria were stained in live cells using the potentiometric dye, tetramethylrhodamine methyl ester (TMRM, 20nM, T668, ThermoFisher Scientific, Burlington, ON, Canada) and imaged using a Leica SP8 confocal microscope (Leica Microsystems Inc., Concord, ON, Canada). Mitochondrial fragmentation was quantified using two reported metrics: mitochondrial fragmentation count (MFC) and a machine-learning based algorithm. Briefly MFC is defined as the total mitochondrial number divided by total mitochondrial area in an image; whereas a machine learning algorithm defines the percentage area of punctate, intermediate and filamentous mitochondria in an image, as previously reported(1).

##### 1.4. Immunofluorescence imaging of endogenous Drp1

For the mitochondrial Drp1 localization studies in fixed cells (representing endogenous Drp1), PAH PASMC were grown to 60-80% confluency, washed with PBS, and treated with 0.1μM Drpitor1a for 5 hours (or DMSO control). During the last 45 minutes of Drptor1a incubation, cells are treated with MitoTracker Deep Red (MTDR) in 37 °C. Cells were then fixed using paraformaldehyde 4% and incubated for 15 minutes at 37 °C. Cells were then permeabilized using 0.5% triton X-100 PBS for 10 minutes rocking at room temperature and subsequently blocked with 2% BSA in PBS with Tween-20 (0.05%) for 35 minutes at room temperature. Primary antibody against Drp1 (#NB110-55237, Novus Biologicals) was diluted (1:400) in 2% BSA PBS with Tween (0.05%) and incubated overnight at 4 °C. Secondary antibody (Anti-rabbit Alexa Fluor 488: #A-11034, ThermoFisher, San Jose, CA) was diluted (1:400) in 2% BSA PBS with 0.05% Tween and incubated for one hour at room temperature covered with aluminum foil to prevent UV exposure. Cells were then mounted onto coverslips using ProLong™ Diamond Antifade Mountant with DAPI (P36962, ThermoFisher Scientific, San Jose, CA). Fixed cells were imaged using a confocal microscope (Leica TCS SP8 Confocal, Leica microsystems Inc., Concord, ON, Canada) using a 1.4 NA 63× Oil objective, pinhole 1 AU, and 2.5× digital zoom. Images were collected at a 512×512-pixel format, and Ex/Em of MTDR was achieved using the WLL laser at 644nm/660nm-700nm on a HyD detector. Excitation and emission of Drp1 antibody was achieved using the WLL laser at 491nm/515nm-535 nm on a HyD detector. All images taken by the microscopist were done with blinding to the treatment.

#### 1.5. Immunoblots

Total cell protein was extracted using cell lysis buffer (9803, Cell Signaling Technology). Cell lysates (40-60 μg) were electrophoresed on a 4%-12% Bis-Tris gel (NP0321BOX, Thermo Fisher

Scientific, Carlsbad, CA, USA), and the proteins were electrotransferred to a 0.2 µm PVDF membrane (LC2002, Life Technologies, Carlsbad, CA, USA). The membrane was then blocked with 5% milk or 5% BSA, followed by overnight incubation with the primary antibody. HRP-linked secondary antibody (NA934V or NA931V, Cytiva, Oakville, ON, Canada) was then added and incubated at room temperature for 1 hour. The detection of specific proteins was carried out using the Amersham ECL Prime Western Blotting Detection Reagent (RPN2236, Cytiva, Buckinghamshire, UK). The following primary antibodies were used: Drp1 (611112, BD Transduction Laboratories, San Jose, CA, USA), Bak (6947, Cell Signaling Technology, Danvers, MA, USA) and Bcl-2 (2872, Cell Signaling Technology, Danvers, MA, USA), β-actin (A5441, Sigma-Aldrich, St. Louis, MO, USA).

##### 1.6. Cell proliferation

hPASMC proliferation was measured 48 hours post-Drpitor1a treatment (1 µM) using a Click-iT EdU Alexa Fluor 488 Imaging Kit (C10420, ThermoFisher Scientific, Burlington, ON, Canada), as previously described (and see on-line supplement) (2). EdU was allowed to incorporate for 16 hours before flow cytometry analysis. The percentage of cells incorporated with EdU was determined using a SONY SH800S cell sorter (SONY Biotechnology, San Jose, CA, USA) and data were analyzed using FlowJo software (Becton, Dickinson & Company, Franklin Lakes, NJ, USA).

##### 1.7. Cell apoptosis

hPASMC apoptosis was determined by the Annexin V+ pro-pidium iodide (PI) method, using Alexa Fluor 488 Annexin V/Dead Cell Apoptosis kit (V13241, ThermoFisher Scientific,

Burlington, ON, Canada). Cells were treated with Drpitor1a (5 μM) and collected 48 hours after compound treatment. The percentage of Annexin V (+) and PI (+) cells was determined using the SONY SH800S flow cytometer.

#### 2. *In vivo* studies

The animal experiment protocol was reviewed and approved by the University Animal Care Committee of Queen's University (protocol number: 2021-2112). All animals received humane care in compliance with the 'Principles of Laboratory Animal Care' formulated by the National Society for Medical Research and the 'Guide for the Care and Use of Laboratory Animals' prepared by the Institute of Laboratory Animal Resources and published by the National Institutes of Health (NIH Publication No. 86-23, revised 1996). Careful attention was paid to adhering to best scientific practices, including the use of both sexes of animals, randomization, blinding of investigators and use of adequate sample size, as previously described(3).

##### 2.1. Pharmacokinetics analysis:

Male and female Sprague-Dawley (SD) rats weighed 250~350g (Charles River, NY, USA) (n=3/sex) were used for the pharmacokinetics study of Drpitor1a. Study objectives. The main objective of this study was to quantify compound exposure levels of Drpitor1a in the plasma of rats following intravenous or oral administration. Quantification was performed using the selective MRM mode on a short LC column using a fast gradient**.**

**Catheter implantation for pharmacokinetics**

A 3 Fr polyurethane catheter (C30PU-RJV1930, Instech Laboratories Inc., Plymouth Meeting, PA, USA) was implanted in the right jugular vein in anesthetized rats before drug administration (IV) or blood sample collection. Anesthesia was induced with 5% isoflurane and maintained with 1.5~2% isoflurane.

**Drug administration**

For pharmacokinetic studies, rats were allowed to recover for 24 hours post-catheterization and then a single dose of Drpitor1a (1mg/kg or 5mg/kg, dissolved in DMSO) was administered IV through the jugular vein catheter or PO by oral gavage without the use of anesthesia.

***in vivo* samples**

Plasma samples were collected at t= 0, 5’, 30’, 1h, 2h, 4h, 6h, 8h and 24h from the jugular vein catheter after intravenous or oral administration of 1 or 5 mg/kg in male or female rats (N=3). The rats were not under anesthesia during blood sampling. Plasma samples were kept frozen at -80°C. Blank $K_2EDTA$ SD rat plasma was obtained from Bioreclamation IVT.

**Test Compounds**

Stock solutions of Drpitor1a was prepared from powder at 10 mM in DMSO and was used for the preparation of calibration curve and QCs.

**Equipment**

Sorvall centrifuge (model RT6000D, Fisher Scientific, Ottawa, ON, Canada). LC/MS/MS AB/SCIEX 4000 QTRAP (Agilent 1100 series HPLC system consisting of autosampler G1367A, column heater G1316A and binary pump G1312A).

**Pharmacokinetic plasma samples extraction**

The plasma tubes were thawed on ice and kept on ice during the preparation. The tubes were mixed and 30 µL of plasma samples were transferred in a 96 wells plate. 75 µL of precipitation solution were added (80/20 acetonitrile/methanol, containing 0.1 µM of loperamide as the internal standard). After mixing, the plate was centrifuged for 15 min at 4000 rpm, 4°C. 30 µL of supernatant were transferred into an HPLC plate, and 2 volumes (60 µL) of water + 0.1% formic acid were added.

**Pharmacokinetics calibration curves preparation**

16-point calibration curve was prepared in blank SD rat plasma, ranging from 0.002 to 15 µM. Stock solutions of Drpitor1a was prepared in DMSO at 10 mM and was diluted to 1 mM in 1:1 methanol/acetonitrile. Standards in 1:1 methanol/acetonitrile were prepared at 750, 500, 250 and 100 µM and QC at 150 µM. Serial dilutions 1:10 were performed in solvent and 1 µL of standard solutions were spiked in 49 µL of blank plasma (total volume 50 □L). Curve points: 0.002, 0.005, 0.01, 0.015, 0.02, 0.05, 0.1, 0.15, 0.2, 0.5, 1, 1.5, 2, 5, 10 and 15 µM. QC points: 0.003, 0.03, 0.3 and 3 µM.

**Pharmacokinetics bioanalysis**

Samples were analyzed in MRM mode using LC-MS/MS. The LC-MS/MS conditions are summarized in Table E1**.** The calibration curves were plotted using the ratio of analyte peak area and IS peak area, using a quadratic regression. PK parameters were calculated using Kinetica Software for PK/PD Data Analysis, Simulation and Reporting (Thermo Fisher), based on the plasmatic concentrations of each animal.

**Table E1.** Summary of LC-MS/MS conditions for Drpitor1a

| | | | | |
|---|---|---|---|---|
| **HPLC** | Agilent 1100 Series | | | |
| **MS/MS** | AB/SCIEX 4000 QTRAP | | | |
| **Software** | Analyst 1.6.2 | | | |
| **Ionisation mode** | Turbo Electrospray | | | |
| **Scan mode** | Multiple reaction monitoring (MRM) using 60 msec dwell time, positive mode | | | |
| **Analytes parameters** | **Compound** | **MRM** | **Declustering (V)** | **CE (V)** |
| | Drpitor1a | 249.2 > 221.2 | 100 | 40 |
| | Loperamide | 477.1 > 266.0 | 70 | 36 |

| | | |
|---|---|---|
| **Source parameters** | Gas Temp (°C) | 600 |
| | Gas flow 1 and 2 | 50 and 60 |
| | Curtain Gas | 20 |
| | Capillary (V) | 5500 |
| **Column temperature** | 45°C | |
| **Sample temperature** | 5°C | |
| **Injection volume** | 4 μL | |
| **Column** | Kinetex C8, 30 x 3 mm, 2.6 um | |
| **Flow rate** | 0.70 mL/min | |
| **Mobile phase** | A: H2O + 0.1% formic acid<br>B: Acetonitrile + 0.1% formic acid | |
| **Gradient** | 2 to 98% B in 1.5 min, plateau at 98% B for 1 min | |

#### 2.2. Tissue concentration analysis

Right ventricle and lung samples were collected at 24 hours post administration of a single dose of Drpitor1a from the rats that underwent pharmacokinetics study. Tissue samples were snap frozen and stored at -80 °C. Tissue concentration of Drpitor1a was measured using HPLC coupled with mass spectrometry at University of Alberta.

**Tissue concentration : chemicals and materials**

Ammonium acetate (Optima LC/MS grade), ammonium hydroxide, water and acetonitrile (HPLC/MS grade) were purchased from Fisher Scientific Company (Ottawa, ON, Canada). Drpitor1a was obtained from Dr. Archer's lab.

**Tissue concentration : sample preparation**

800µl of a fresh prepared solvent mixture of methanol and water (80:20) was added to tissue samples. The mixture was votexed and homogenized (10pulses, at 10) using a sonic dismembrator. It was incubated on ice-bath for 20mins, and the mixture was votexed every 5 mins during the incubation, followed by centrifugation at 13,000 xg for 15mins at 4oC. The supernatant was collected and evaporated by speed-Vac. The dried extracts were stored at -80oC and then re-dissolved in 100µl of solvent (acetonitrile:methanol, 50:50) prior to HPLC/MS /MS analysis.

**Tissue concentration : HPLC/MS /MS analysis**

Standard and sample solutions were analyzed using an ARIA MX HPLC system (Thermo Fisher Scientific, San Jose, CA) coupled to Orbitrap LTQ XL Mass spectrometer (Thermo Scientific, San Jose, CA) and using Thermo Xcalibur software for data acquisition and analysis. A Xbridge BEH 150mmx2.1mm Amide column, 2.5μm particle size (Waters, Milford, MA) was employed for LC separations. The column temperature was controlled at 25$^{o}$C. The mobile phase A was acetonitrile and B was water containing 10mM ammonium acetate and 0.1% formic acid. The gradient was as follows 0-0.1min, 5% B 0.1-2min, linear gradient to 15% B 2-3min, linear gradient to 25%B; 3-6 min, linear gradient to 80% B and then back to 5% B at 6.1min and equilibrium for 9min. The flow rate of mobile phase was 200μl/min and the cycle time was 15min/injection. The orbitrap mass spectrometer was operated under electrospray positive ion mode. Ionization voltage was set at +4KV. Nitrogen was used as sheath gas, aux gas and sweep gas. They were set at Sheath gas 35, aux gas 30 and sweep gas 3 (Arbitrary units). The Ion source temperature and capillary temperature were at 350oC and 325oC, respectively. Acquisition was carried out in full scan mode with mass range from 50 to 1000amu with resolving power set to a nominal value of 60,000 at full width half-maximum at m/z 400. Within the same analysis, tandem mass spectrometry (MS/MS) was performed for ion at m/z 249.07 using high energy collision dissociation (HCD) at 65eV at resolution at 30,000. Mass calibration and tuning was done by infusing of LTQ Velos ESI Positive Ion Calibration Solution (Thermo Fisher Scientific, Rockford, IL) prior to performing HPLC/MS/MS analysis.

A five-point calibration curve of Drpitor1a at concentration of 0.25 μg/L, 0.625 μg/L, 1.25 μg/L 6.25 μg/L and 12.5 μg/L was constructed based on the peak area of the Driptor1a versus the Driptor1a concentration. The samples concentration was calculated with this calibration curve.

#### 2.3. Efficacy of Drpitor1a on MCT-PAH

##### 2.3.1. Sex and sample size

**Pilot study:** The sample size in the pilot study was small, n=3 rats/sex in the MCT+NS groups and 4 rats/sex were used in the two MCT+Drpitor1a groups (0.1 mg/kg and 1.0 mg/kg IV every 48 hours, see protocol **Figure 4A**. The pilot study was designed to assess feasibility and tolerability of Drpitor1a and observe potential efficacy signals (defined as a trend to reduce adverse remodeling of small PAs and regress RV hypertrophy in MCT-PAH). It was not powered to observe statistical differences in hemodynamics between the sexes.

**Confirmation study:** Because Drpitor1a lacked benefit in male rats in the pilot study, only female rats were used in the confirmation study. We compared Drpitor1a to vehicle in normal females (n=6 vs 8) and MCT-PAH females (n=9 and 11), respectively. Drpitor1a treatment (1.0 mg/kg every 48 hours) began 17-days post-MCT with echocardiography and cardiac catheterization performed 28-29 days post-MCT. Sample size was based on a power calculation data informed by the pilot study and was sufficient to ensure 80% power to detect a significant Drpitor1a effect with a $p<0.05$.

##### 2.3.2. Animal model of MCT-PAH:

PAH was induced in SD rats (Charles River, NY, USA) weighing 200-250g by a single subcutaneous injection of monocrotaline (MCT, 60mg/kg, C2401, Millipore Sigma, Oakville, ON, Canada), as previously described (4). The same volume of PBS was used as control.

##### 2.3.3. Jugular vein indwelling catheter implantation

On day 15 post-MCT/PBS injection, rats were anesthetized with inhaled isoflurane (5% for induction, 1.5~2% for maintenance) and a 3Fr polyurethane catheter (C30PU-RJV1931, Instech

Laboratories, Inc., Plymouth Meeting, PA) was implanted in the left jugular vein (LJV) for drug infusion. Catheters remained in place for the duration of the study (~2 weeks), and were infused with 0.1 mL of 100U/mL heparin every 24 hours to maintain their patency. Tramadol (0.1mL, 50 mg/mL, SC) and meloxicam (Metacam, 0.1mL, 5 mg/kg, SC) were used from day 0 to day 2 post-operation for pain management.

###### 2.3.4. Drpitor1a treatment of female rats with established MCT-PAH

Drpitor1a or control vehicle was administered IV from the jugular vein, indwelling, catheter every 48 hours from day 17 to day 27. A stock solution of Drpitor1a (5 mg/ml) was prepared in anhydrous DMSO and filtered. The stock solution was further diluted in normal saline (NS) to 5% DMSO (v/v) for infusion. The final dose of Drpitor1a for treatment was 1 mg/kg body weight. The same volume of 5% DMSO was used for the control groups.

###### 2.3.5. Echocardiography

Doppler, 2-dimensional (2-D), and time-motion (M-mode) ultrasound was performed on day 14 and day 28 post-MCT/PBS injection using a high-frequency ultrasound platform (Vevo 2100, Visual Sonics, Toronto, ON, Canada) and a 37 MHz probe, as previously described (4). The following parameters were determined using echocardiography: pulmonary artery acceleration time (PAAT), systolic velocity time integral (VTI) in the main pulmonary artery (PA), main PA inner radius (IR), at the level of the pulmonary outflow tract during mid-systole, the diastolic and systolic thickness of the right ventricular free wall (RVFW), and tricuspid annular plane systolic excursion (TAPSE). Cardiac output (CO) was estimated using the following formula:

$$\mathrm{CO} = \text{Heart rate (HR)} \times \mathrm{VTI} \times \pi \times \mathrm{IR}^2$$

###### 2.3.6. Hemodynamics-left and right heart catheterization

Pulmonary hemodynamics were measured on day 29 by right heart catheterization (RHC) using a 1.9F pressure-volume catheter (FTH-1912B-6018, Transonic Systems Inc., Ithaca, NY, USA) delivered to the RV via a closed-chest approach, as previously described (1). Right ventricular systolic pressure (RVSP) and CO were determined using RHC.

After pulmonary hemodynamics were determined, the chest was opened and left-ventricular end-diastolic pressure (LVEDP) was measured by direct LV puncture. Cardiac index (CI), mean pulmonary artery pressure (mPAP) and pulmonary vascular resistance index (PVR) were calculated using the following formulae:

$$\mathrm{CI} = \frac{\mathrm{Cardiac\ output\ (CO)}}{\mathrm{Body\ weight}}$$

$$\mathrm{mPAP} = 0.61 \times \mathrm{RVSP} + 2\ \mathrm{mmHg}\ (5)$$

$$\mathrm{PVR} = \frac{\mathrm{mPAP} - \mathrm{LVEDP}}{\mathrm{CO}}$$

##### 2.3.7. Blood collection

Whole blood was collected via cardiac puncture on day 29 after RHC. The following tests were performed: complete blood count, chemistry panel and chemistry renal profile analyses (Antech Diagnostics, Inc., Fountain Valley, CA, USA).

**Table E2. Average ± Standard deviation of the pharmacokinetic parameters in the rats receiving oral administration of Drpitor1a.**

| Oral administration | | | | |
|---|---|---|---|---|
| **Parameters** | **Male - 1mg/kg** | **Female - 1mg/kg** | **Male - 5mg/kg** | **Female - 5mg/kg** |
| AUC0-t (nM*h) | 143.4 ± 106.0 | 162.7 ± 96.1 | 526.4 ± 170.9 | 1044.9 ± 406.2 |
| AUC0-∞ (nM*h) | 165.7 | 246.7 ± 95.4 | 1006.8 ± 113.4 | 1448.0 |
| MRT (h) [(a)] | 8.8 | 4.4 ± 0.4 | 10.9 ± 3.0 | 6.9 |
| kel (h) | 0.1 | 0.2 ± 0.01 | 0.1 ± 0.07 | 0.2 |
| t1/2 terminal (h) | 5.5 | 3.2 ± 0.2 | 6.6 ± 2.8 | 4.0 |
| Tmax (h) | 2.5 ± 3.0 | 3.0 ± 2.7 | 3.4 ± 3.0 | 3.3 ± 4.2 |
| Cmax (nM) | 42.2 ± 33.8 | 48.6 ± 32.3 | 87.7 ± 32.5 | 141.4 ± 46.7 |
| CLp (mL/min) | 132.6 | 72.5 ± 29.0 | 100.4 ± 9.4 | 53.6 |
| Vd (L) | 48.7 ± 24.3 | 19.4 ± 9.3 | 66.8 ± 22.4 | 41.2 ± 26.8 |

(a) MRT = time for elimination of 63.2% of the dose.
AUC: Area Under the Concentration-Time Curve
MRT: Mean Residence Time
kel: elimination rate constant
t1/2: terminal half-life
Tmax: the time it takes for Drpitor1a to reach its maximum concentration in the bloodstream after administration
Cmax: the maximum concentration of Drpitor1a in the bloodstream after it has been administered
CLp: the clearance of Drpitor1a from the plasma
Vd: the volume of distribution

References:

1. Chen KH, Dasgupta A, Lin J, Potus F, Bonnet S, Iremonger J, Fu J, Mewburn J, Wu D, Dunham-Snary K, Theilmann AL, Jing ZC, Hindmarch C, Ormiston ML, Lawrie A, Archer SL. Epigenetic Dysregulation of the Dynamin-Related Protein 1 Binding Partners MiD49 and MiD51 Increases Mitotic Mitochondrial Fission and Promotes Pulmonary Arterial Hypertension: Mechanistic and Therapeutic Implications. *Circulation* 2018; 138: 287-304.
2. Wu D, Dasgupta A, Chen KH, Neuber-Hess M, Patel J, Hurst TE, Mewburn JD, Lima PDA, Alizadeh E, Martin A, Wells M, Snieckus V, Archer SL. Identification of novel dynamin-related protein 1 (Drp1) GTPase inhibitors: Therapeutic potential of Drpitor1 and Drpitor1a in cancer and cardiac ischemia-reperfusion injury. *FASEB J* 2020; 34: 1447-1464.
3. Provencher S, Archer SL, Ramirez FD, Hibbert B, Paulin R, Boucherat O, Lacasse Y, Bonnet S. Standards and Methodological Rigor in Pulmonary Arterial Hypertension Preclinical and Translational Research. *Circ Res* 2018; 122: 1021-1032.
4. Tian L, Wu D, Dasgupta A, Chen KH, Mewburn J, Potus F, Lima PDA, Hong Z, Zhao YY, Hindmarch CCT, Kutty S, Provencher S, Bonnet S, Sutendra G, Archer SL. Epigenetic Metabolic Reprogramming of Right Ventricular Fibroblasts in Pulmonary Arterial Hypertension: A Pyruvate Dehydrogenase Kinase-Dependent Shift in Mitochondrial Metabolism Promotes Right Ventricular Fibrosis. *Circ Res* 2020; 126: 1723-1745.
5. Chemla D, Castelain V, Provencher S, Humbert M, Simonneau G, Herve P. Evaluation of various empirical formulas for estimating mean pulmonary artery pressure by using systolic pulmonary artery pressure in adults. *Chest* 2009; 135: 760-768.
